# Multimodal profiling unveils a reversible breast basal-like cell state in AKT-inhibitor resistant tumours

**DOI:** 10.1101/2025.06.06.658331

**Authors:** Colin D.H. Ratcliffe, Hugh Sparks, Giulia L.M. Boezio, Yuriy Alexandrov, Robert P. Jenkins, Alix Le Marois, Pablo Soro-Barrio, Fatemeh Ahmadi Moughari, Heather Hulme, Rebecca J. Lee, Stephanie Strohbuecker, Sudeep Joshi, James K. Ellis, Giacomo Fabrini, Lucy Flint, Anne-Marie Fortier, Nils Gustafsson, Ana C. Cunha, Louis Tully, Morag Park, Fabian Fröhlich, James I. MacRae, Rachael Natrajan, Andrea Serio, James Briscoe, Andy Riddell, Simon T. Barry, Christopher Dunsby, Erik Sahai

**Affiliations:** Tumour Cell Biology Laboratory, The Francis Crick Institute, London, NW1 1AT, UK; Photonics Group, Imperial College London, South Kensington Campus, London, SW7 2AZ, UK; Developmental Dynamics Laboratory, The Francis Crick Institute, London, NW1 1AT, UK; Bioinformatics & Biostatistics, The Francis Crick Institute, London, NW1 1AT, UK; The Breast Cancer Now Toby Robins Research Centre, The Institute of Cancer Research, London SW3 6JB, UK; Integrated Bioanalysis, Clinical Pharmacology and Safety Sciences, AstraZeneca R&D, Cambridge, CB2 0AA, UK; Neural Circuit Bioengineering and Disease Modelling Laboratory, The Francis Crick Institute, London, NW1 1AT, UK; Basic & Clinical Neuroscience Department, Institute of Psychiatry, Psychology and Neuroscience (IoPPN), King’s College London, Maurice Wohl Clinical Neuroscience Institute, London, UK; Metabolomics, The Francis Crick Institute, London, NW1 1AT, UK; Dynamics of Living Systems Laboratory, The Francis Crick Institute, London, NW1 1AT, UK; Goodman Cancer Institute, McGill University, Depts Biochemistry and Oncology, McGill University, Montréal, Canada; Viral Vector Core, The Francis Crick Institute, London, NW1 1AT, UK; Dementia Research Institute (UK DRI), London, United Kingdom; Flow Cytometry, The Francis Crick Institute, London, NW1 1AT, UK; OTD, Early Oncology, AstraZeneca, Cambridge, CB2 0AA, UK

## Abstract

The PI3K/AKT/mTOR pathway is central to cell metabolism and growth. However, pharmacological inhibition of the pathway is not uniformly effective across cancer types, or even within a single cancer model. In this study, we leverage oblique plane microscopy of triple negative breast cancer organoids, as well as lineage tracing to uncover a source of heterogeneity. Non-genetic resistance to AKT inhibition is associated with basal cell features of normal breast epithelium and the master transcription factor of basal cell state, ΔNp63α, is sufficient to confer resistance. Cells can transition between states within four weeks and therefore, AKT inhibition only delays tumour growth, with tumours rich in KRT14+ cells resulting. Thus, under selection, triple negative breast cancer exploits a repertoire of cell states inherent to the breast.

## Introduction

Heterogeneity between cancer cells can undermine the efficacy of therapy. In some cases, resistant cells arise after exposure to therapy, with cells entering a persister cell state prior to transitioning to a resistant state (*1, 2*). In other cases, tumours contain a subset of cells that are therapy resistant before treatment. In both, the cause of resistance can be genetic or non-genetic (*3–5*).

The PI3K pathway is activated in the majority of cancers, with mutations in *PIK3CA*, *AKT*, and *PTEN* observed in over half of all solid tumours, leading to increased glucose uptake as well as increased mTOR, and S6 kinase activity (*6–11*). These changes ultimately lead to increased glycolysis and protein synthesis, enabling tumour growth (*12–18*). Pharmacological inhibition of either PI3K or AKT provides clinical benefit in certain contexts, although durable responses are rare with reactivation of mTOR and S6 kinase frequently observed (*19–25*). AKT inhibition is a novel FDA-approved strategy to treat advanced ESR1+ breast cancer with abnormal *PIK3CA*, *AKT1* or *PTEN* (*26*), but does not provide overall survival benefit in triple-negative disease (TNBC) despite frequent mutational activation of PI3 kinase signalling. Therefore, mutations alone are not sufficient to explain tumour response or resistance. Recent work has suggested that tumour regression through AKT inhibition requires mechanisms relating to involution, supporting a role for non-genetic heterogeneity in cell states for resistance to AKT inhibitors (*27*).

By using patient-derived organoids, we identify a subpopulation that exhibits a signature positive for KRT14, a marker of basal cells in the normal breast, and resists a bottleneck selection pressure imposed by AKT kinase inhibition. The resistant cell state is reversible and can be driven by ΔNp63α overexpression. Therefore, we propose that resistance to AKT inhibition in TNBC exploits a repertoire of cell states and reversible switching mechanisms inherent to the biology of the breast.

## Results

### Establishment of a TNBC model recapitulating tumour heterogeneity

To investigate the phenotypic heterogeneity of triple negative breast cancer, we employed patient-derived xenograft organoids (PDXOs). GCRC1915, recapitulates histological and molecular features frequently observed in TNBC, including genetic alterations such as *TP53* mutation, chromosomal gain of *PIK3CA* and low expression of PTEN (*28*). GCRC1915 organoids were dissociated and transduced as single cells with the LARRY barcode library (*29*). After seven days, GFP^+^ organoids were enriched using a custom-built organoid sampling platform coupled to live-cell microscopy, termed VISIBLE (*30*), and injected into mice (Fig. S1A – left branch). After five weeks, tumours were harvested and single cell RNA sequencing of human cells was performed. For brevity, we termed this framework **L**arry **B**arcoding **E**xpansion and **T**reatment with **O**rganoid **S**orting and **S**equencing (LBETROSS) (Fig S1A).

We identified three clusters by unsupervised detection (Clusters I, II and III) and these were enriched for specific gene modules (GMs) previously described in human breast cancer (*31*) (Fig. 1A & Fig. S1B). All GMs, except GM5, were identified in our model (Fig. S1C). Barcode analysis identified 60 clonal groups with 4 or more cells (Fig. 1B). Intriguingly, clonal groups were scattered between clusters with binomial testing revealing that 95% of these were present in multiple clusters (Fig. 1C & D). These data demonstrate that our model recapitulates non-genetic transcriptional heterogeneity observed in human patients, and that cells transition between transcriptional states within a six-week timeframe.

**Fig. 1.**
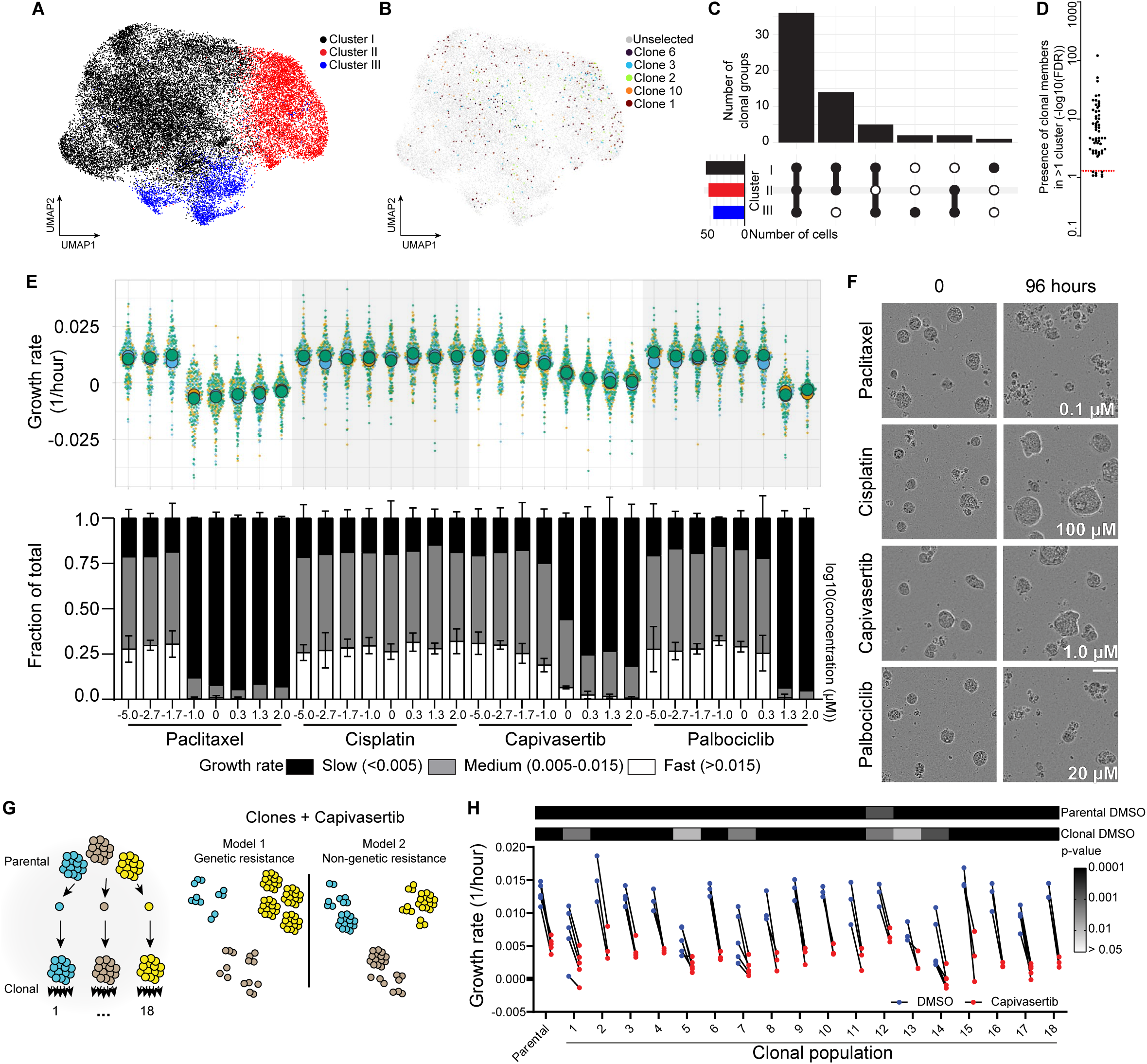
Non-genetic heterogeneity underpins response to Akt kinase inhibition. UMAP (uniform manifold approximation and projection) representation of GCRC1915 patient-derived xenograft single-cell RNA sequencing datasets with colours indicating **(A)** cluster membership (resolution = 0.1) or **(B)** LARRY barcode identity of the 5 largest clonal groups. **(C)** Upset plot displaying cluster membership of clonal groups **(D)** Plot of the results from a binomial test to evaluate the presence of clonal group members within more than one cluster. **(E)** Growth rates of organoids treated with indicated drug conditions for 4 days. In the upper panel, data are represented as Superplots where dots represent organoids (n=8682), circles represent experimental means (n=3) and colours indicate experiment identity. In the lower panel, relative fractions of organoid growth rates for each condition is shown. **(F)** Representative brightfield micrographs of GCRC1915 PDXOs. Scale, 50 μm. **(G)** Schematic outline of organoids where colour represents clonal identity, followed by two possible experimental outcomes. **(H)** Paired mean growth rates of GCRC1915 PDXO clones treated with DMSO or Capivasertib (n=3 or 5) with calculated p-values (Holm-Šídák’s multiple comparison test).

### Non-genetic heterogeneity in response to AKT inhibition

Next, we investigated the relevance of pre-existing heterogeneity on treatment response using timelapse imaging, which provides information about organoid size at the start of treatment and subsequent growth rate. Organoids were treated with increasing concentrations of the AKT inhibitor – Capivasertib, two conventional chemotherapies – paclitaxel and cisplatin, and the CDK4/6 inhibitor Palbociclib. Resistance to Cisplatin, observed in the GCRC1915 PDX (*28*), was recapitulated by the organoid model, along with sensitivity to Paclitaxel. The organoids were also sensitive to Palbociclib. Intriguingly, response to Capivasertib diverged from those of other cytotoxic drugs (Fig. 1E). In particular, the Capivasertib dose-response had a shallower slope and there was a greater proportion of organoids that maintained a moderate growth rate even at high doses of the drug. This heterogeneity was clearly evident in timelapse movies, with some organoids continuing to grow while others of similar size and in the same field of view did not (Fig. 1F, Fig. S2A and Movie S1). Comparable results were obtained with the AKT inhibitor Ipatasertib (Fig. S2A, B & C). Analysis using inhibitors of PI3K, mTOR, and Cytohesin suggested that PIK3CA was likely to be the key isoform acting upstream of AKT, and that mTOR was a critical downstream effector (Fig. S2D).

The heterogeneous response to AKTi indicated a pre-existing resistant population of cells within the PDXO culture. To explore if this might be underpinned by a genetic subclone, we isolated single cell-derived sub-lines (Fig. 1G). A power calculation based on the 17% of organoids that continued growing in the presence of Capivasertib indicated that 18 sub-lines would be sufficient to determine if a pre-existing resistant sub-clone was present with a significance level of <0.05. Notably, all 18 sub-lines were sensitive to Capivasertib and none grew in the presence of Capivasertib at the same rate as DMSO-treated controls, arguing against a genetic mechanism of resistance (Fig. 1H).

### Differential dynamics of AKT regulation and coupling to glucose distinguish organoids resistant to AKT inhibition

The failure to respond to a therapy could reflect ineffective inhibition of the target, the lack of coupling between the target and key biological processes, or the lack of dependence of the resistant cells on the biological process regulated by the drug target. We sought to resolve these differing scenarios by measuring both single-cell AKT activity and glucose metabolism, along with whole-organoid features using fluorescent biosensors combined with a custom-built dual-view oblique plane microscope (dOPM). Methodological development and benchmarking are discussed in supplemental text and Fig S3 & 4. Biosensor expressing organoids were imaged immediately prior to Capivasertib administration, 1 day (D1) and 4 days (D4) later. To summarise our observations, we plotted the data where each column represents an organoid, and the columns are ranked based on growth rate (Fig. 2A shown in the upper panel). The central and lower panels indicate the glucose sensor signal and AKT activity at day 4, respectively. Every cell in an organoid is represented as a thin horizontal bar contributing to the height of the column, with taller columns indicating more cells in the organoid, and the cells are ranked in order of increasing glucose sensor FRET signal. These analyses reveal the relationships between AKT, glucose levels, and proliferation in the presence or absence of AKT inhibition (Fig. 2B).

**Fig. 2.**
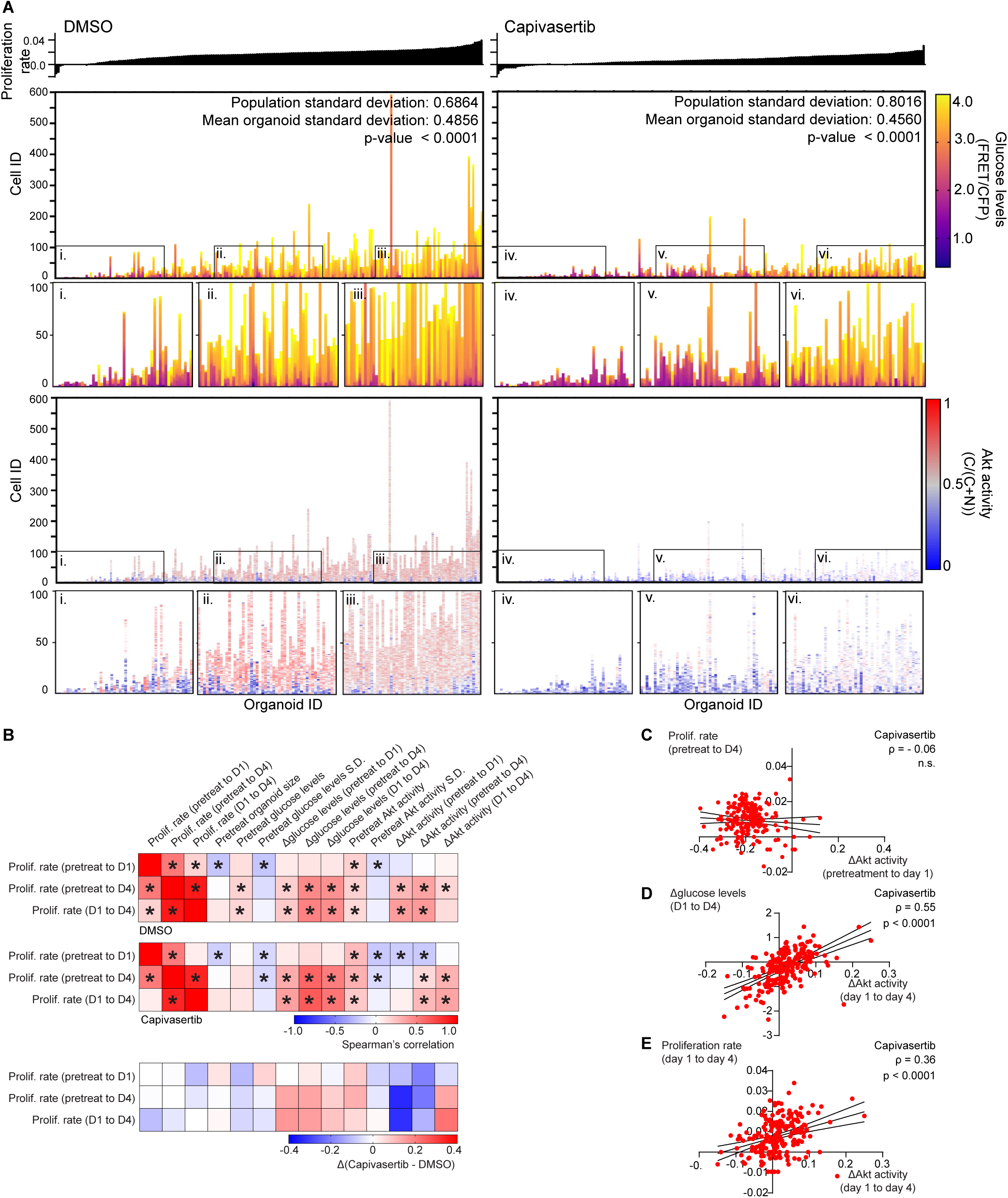
Collective differences to Akt kinase inhibition underpin responses to Capivasertib. **(A)** Heatmap representation of organoids. Each column represents an organoid and each row represents a cell from that organoid. Growth rates for organoids are plotted above and pseudo-colour represents single cell FRET or Akt activity. Organoids are arranged in order of increasing growth rate. Cells are arranged in order of increasing glucose FRET sensor signal. Insets encompass the bottom, middle and top 50 organoids in the dataset **(B)** Heatmap representation of (top and middle) Spearman’s correlation coefficients between OPM-derived metrics and (bottom) the difference between Capivasertib and DMSO-treated conditions. * p < 0.05 two-tailed **(C)** Plotted proliferation rates of organoids with 1 μM Capivasertib as a function of the change in Akt values at day 1 relative to pretreatment values. **(D)** Plotted change in Glucose values at day 4 relative to day 1 of organoids with 1 μM Capivasertib as a function of the change in Akt values at day 4 relative to day 1. **(E)** Plotted proliferation rate from day 1 to day 4 of organoids with 1 μM Capivasertib as a function of the change in Akt values at day 4 relative to day 1.

Firstly, as expected, DMSO-treated organoids with higher baseline AKT signalling and glucose levels grew slightly faster; and AKT inhibition reduced AKT activity and glucose levels (Fig. 2A). The largely monochromatic columns indicated that cells within an organoid respond similarly. This was confirmed by the narrower standard deviation of FRET signal within organoids (DMSO-0.4856 and Capivasertib-0.4560), compared to the overall population (DMSO-0.6864 and Capivasertib-0.8016). However, not all organoids responded in the same way, with the fastest 15% Capivasertib-treated organoids continuing to grow faster than the DMSO median (0.018 1/hour), and these maintained their glucose levels (Fig. 2A.vi).

Secondly, the observed heterogeneous response is not due to variability in Capivasertib targeting since the observed proliferation rate following inhibition did not correlate with the reduction in AKT activity 1 day after drug treatment (Fig. 2C). Pre-treatment features, such as glucose levels or size, did not predict coupling between AKT signalling and resistance (Fig. S5A & B).

Finally, longitudinal tracking of organoids to 4 days enabled us to explore the relationship between the durability of response to Capivasertib and the efficacy of the drug in stopping growth. Organoids that showed an increase in AKT activity between 24 - 96 hours had an increase in glucose levels during the same time window (Fig. 2D). Crucially, these were the organoids that continued to grow in the presence of drug (Fig. 2E). These data establish a clear link between heterogeneity in the durability of AKT inhibition and the downstream consequences of glucose regulation and cell growth. These data argue for inter-organoid heterogeneity in the wiring of AKT signalling feedback, with AKTi resistant organoids having a faster restoration of glucose levels following AKT inhibition.

### Drug refractory organoids arise from a pre-existing cell state

Diverse drug responses between organoids might reflect heterogeneity in cell states. To address this, we leveraged the LBETROSS pipeline (Figure S1A – right branch). Following barcoding and treatment, we sorted GFP^+^ organoids and performed single cell RNA sequencing (Fig. S6A, B and C). Flow cell imaging and subsequent size analysis confirmed our ability to sort intact organoids from both DMSO and Capivasertib-treated conditions (Fig. S6D).

UMAP analysis of 6623 cells revealed three major clusters, numbered 1-3, and two minor clusters, numbered 4 & 5 (Fig. 3A). All clusters were represented in both control DMSO and Capivasertib-treated conditions, indicating that cell states resistant to drug are present in the drug naïve context (Fig. 3B). Analysis of cell cycle phase revealed that clusters 1, 4, and 5 had a higher proportion of cells in G1 following AKT inhibition, whereas clusters 2 and 3 did not (Fig. 3C). Additionally, clusters 2 and 3 made up an increased proportion of total cells under Capivasertib-treated conditions, supporting a model where clusters 2 and 3 represent drug-resistant populations. Of note, these clusters were enriched for genes associated with basal-like breast cancer and mammary stem cells, including *SOX2, CD44, KRT14, KRT5* and *TP63* (Fig. 3D).

**Fig. 3.**
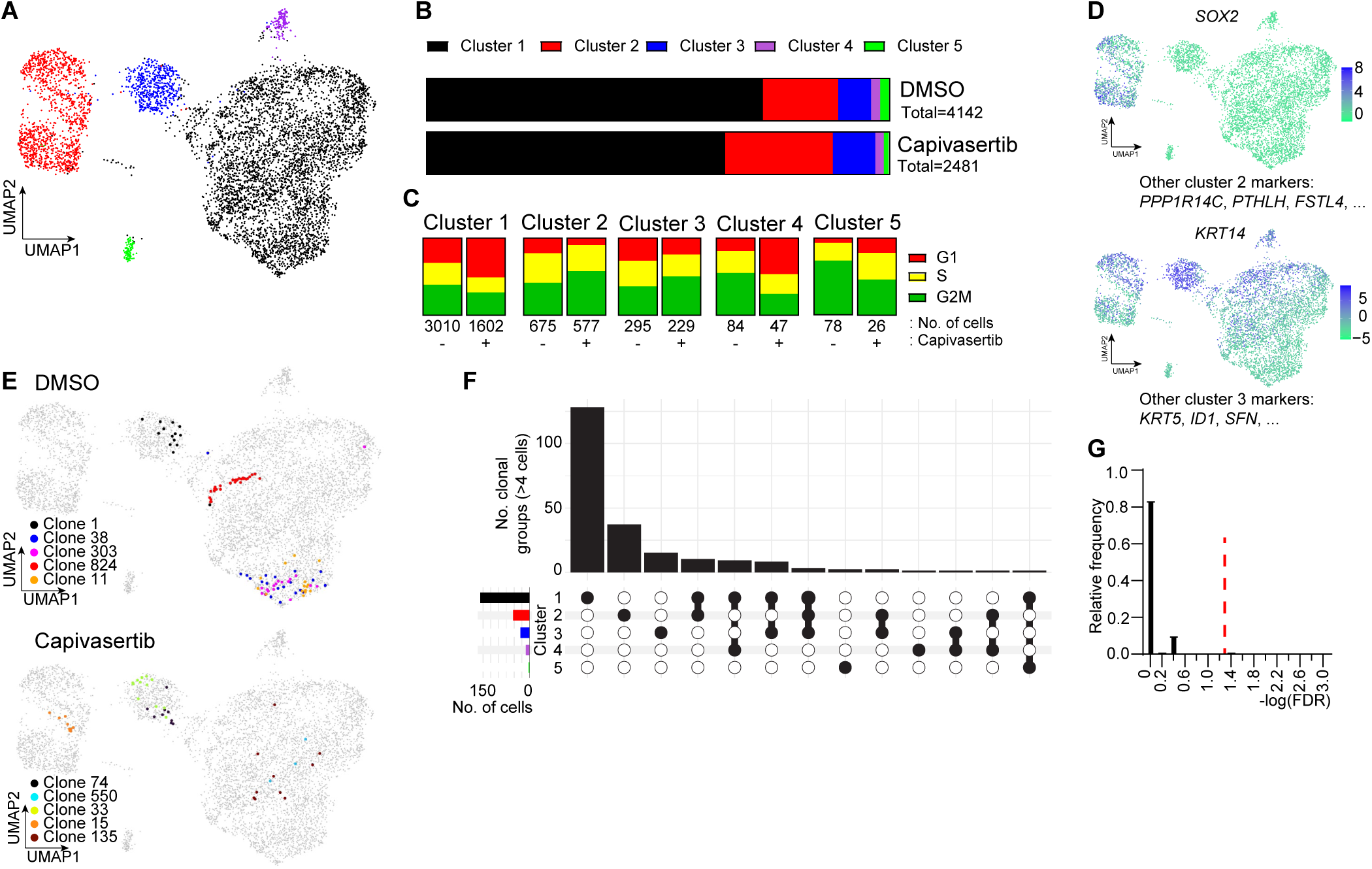
Lineage tracing unveils inter-organoid differences in gene expression. **(A)** UMAP representation of LARRY-barcoded GCRC1915 PDXO cells treated with DMSO or Capivasertib. **(B)** Bar chart depicting the fraction of all cells made up by each cluster defined by gene expression. **(C)** UMAP representation of LARRY-barcoded GCRC1915 PDXO cells with colours indicating gene expression levels **(D)** Bar chart depicting the fraction of total cells in each cell cycle state split by treatment groups (DMSO or Capivasertib) and cluster. **(E)** UMAP representation of LARRY-barcoded GCRC1915 PDXO cells with colours indicating gene barcode identity of the 5 largest clonal groups. **(F)** Upset plot displaying cluster membership of clonal groups. **(G)** Histogram displaying the relative frequencies of the false discovery rates (FDR) for the presence of clonal group members within more than one cluster.

Lineage analysis was performed to identify cell states that contribute to the make-up of a given organoid. Barcodes could be detected in 37-54% of cells and a mean of 314 clonal identities could be detected per sample, with clone sizes ranging from 1 to 29 cells. We restricted our analysis to clonal groups that contained at least 4 cells. Mapping of a barcode to a single transcriptional cluster would indicate no transitions between states occurring during the seven days between barcoding and organoid harvesting. Figure 3E shows the five largest clonal groups from each condition overlayed onto UMAP space where we find that differences in gene expression within a given organoid is limited, with almost no barcodes being present in more than one cluster. Indeed, single cluster-contained clonal groups are the three most common distributions of clonal groups amongst gene expression clusters and binomial testing reveals that >96% of clonal groups confidently belong to a single cluster (Fig. 3F & G). Therefore, within the time frame of the experiment, cells do not transition between states and, consequently, heterogeneity in Capivasertib response between organoids reflects heterogeneity in cell state when the organoid was seeded.

### Long-term resistance to AKT inhibition is linked to a basal cell state

We next set out to determine traits associated with long-term resistance to AKTi. We generated a population of uniformly resistant organoids, treating and passaging them with Capivasertib for four weeks (Fig. 4A). Metabolomic characterisation confirmed that AKTi had the expected effect on control organoids, lowering glycolysis. In contrast, AKTi had a greatly reduced impact on glucose-derived pyruvate and lactate in the resistant organoids (Fig. S7A). These organoids also had increased glucose-derived ^13^C into the TCA cycle intermediates citrate, glutamate, fumarate, aspartate and malate. These data are consistent with Figure 2, where organoids capable of growth under Capivasertib treatment maintain their glucose levels and have a metabolic profile not dissimilar to drug-naïve cells.

**Fig. 4.**
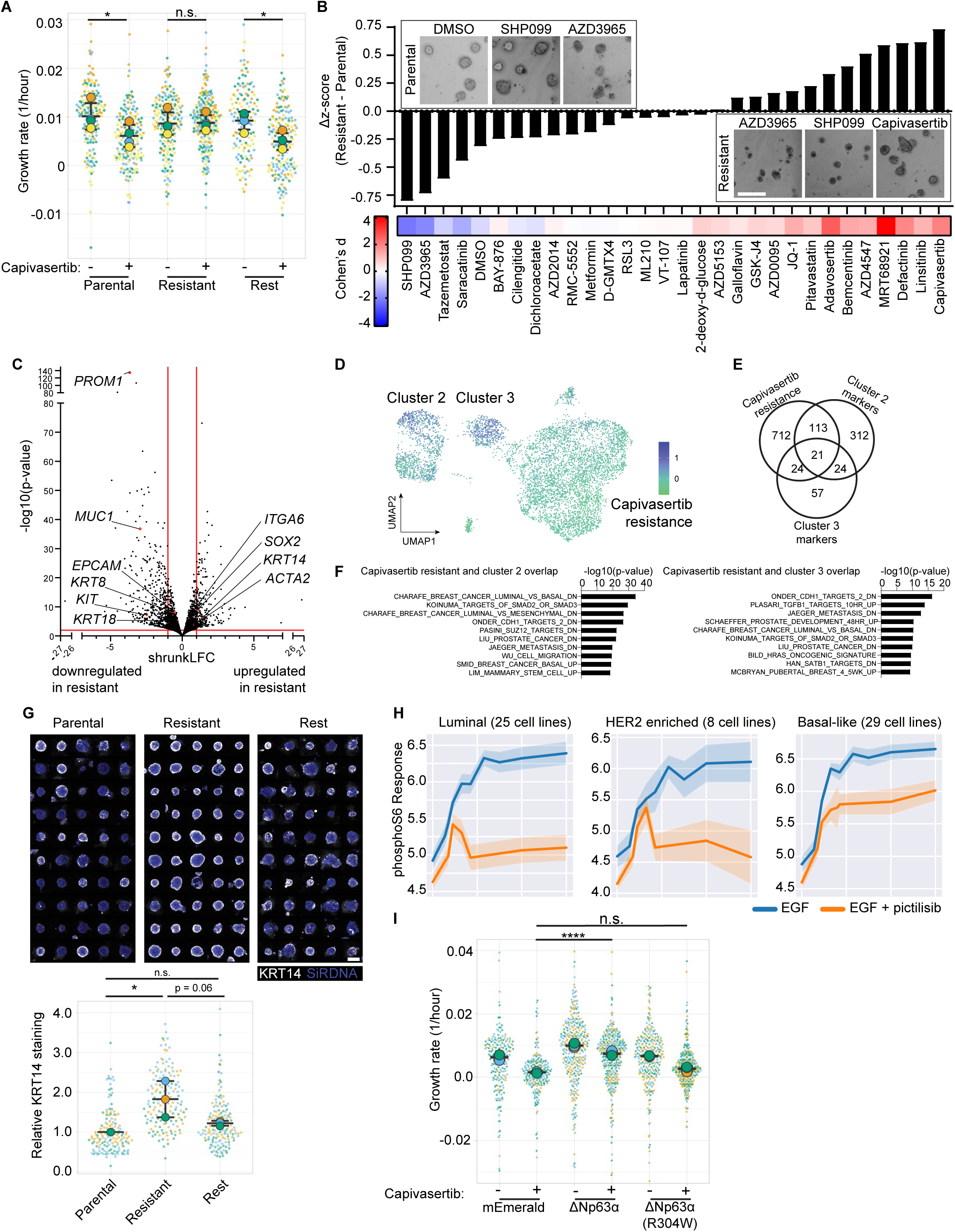
p63 confers resistance to Capivasertib. **(A)** Growth rates of organoids treated with DMSO or Capivasertib for 4 days. Data are represented as Superplots where dots represent organoids (n=570), circles represent experimental means (n=4) and colours indicate experiment identity. **(B)** Bar plot of the difference in z-scores between parents and Capivasertib-resistant organoids treated with the indicated drugs. Heatmap denotes the effect size as measured by Cohen’s d. (n=3) Inset – Representative minimum intensity projections Scale, 200 μm. **(C)** Volcano plot depicting differential gene expression of Capivasertib resistant organoids relative to DMSO controls. **(D)** UMAP representation of LARRY-barcoded GCRC1915 PDXO cells with colours indicating upregulation of Capivasertib resistance gene signature defined by the top 200 upregulated genes identified in Capivasertib-resistant organoids. **(E)** Venn diagram depicting the number and overlap of genes upregulated in Capivasertib-resistant organoids, as well as markers of Clusters 2 and 3 depicted in Fig. 4A. **(F)** Barplots depicting gene set enrichment signature p-values for overlapping between datasets. **(G)** Digital montage of maximum project images of pseudo-coloured micrographs representing DMSO Capivasertib-resistant and Capivasertib-rested organoid lines and associated quantification of normalised KRT14 staining intensity. Data are represented as Superplots where dots represent organoids (n=450), circles represent experimental means (n=3) and colours indicate experiment identity. **(H)** Growth rates of organoids treated with DMSO or Capivasertib for 4 days. Data are represented as Superplots where dots represent organoids (n=1616), circles represent experimental means (n=3) and colours indicate experiment identity. * p < 0.05 **** p <0.0001 (Holm-Šídák’s multiple comparison test).

Despite this, the organoids had not entered a broadly drug-resistant cell state since sensitivity to Paclitaxel was not affected (Fig. S8A & B). We explored this more generally by comparing organoid growth under treatment of a selection of drugs including inhibitors of epigenetic modifiers, tyrosine kinases, integrin signalling, metabolism or activators of ferroptosis (Fig. S8C & D). Supporting a metabolic requirement for glucose, both 2-deoxy-glucose and the Glut1 inhibitor, BAY-876, abolished growth of both parental and resistant organoids. Interestingly, Capivasertib-resistant organoids were no longer sensitive to ULK1 (MRT68921) inhibition (Fig. S8D). However, Capivasertib-resistant organoids were not broadly drug resistant, SHP2 (SHP099), as well as MCT1 (AZD3965) inhibition reduced growth to a greater extent in these organoids suggesting that they have specific molecular vulnerabilities (Fig. 4B).

Data presented thus far support a model where resistance to Capivasertib is driven by cell states. Notably, cessation of Capivasertib treatment led to a reversion to drug sensitivity within 4-6 weeks, further emphasising the non-genetic nature of the AKTi-resistant state (Fig. 4A).

To define the Capivasertib-resistant state we analysed organoids by bulk RNA sequencing (Fig. 4C). Genes up-regulated in long-term resistant organoids mapped to clusters 2 and 3 of our single-cell RNA sequencing dataset and there was significant overlap with markers of these clusters (Fig. 4D & E). Moreover, resistant cells were similar to breast basal cells with high mRNA expression of *SOX2*, *KRT5*, *KRT14*, and *ITGA6* (Fig. 4C & F); high protein levels of ITGA6 and KRT14 and low protein levels of EPCAM (Fig. S9A, B & C). Confirming the reversibility of the resistant phenotype, re-emergence of a ITGA6 low population occurred 3 weeks after removal of Capivasertib from resistant organoids (Fig. S9D) and rested organoids displayed ITGA6, KRT14, and EPCAM levels similar to drug naïve organoids. dOPM analysis confirmed that KRT14 staining was associated with organoid identity and that resistant organoids were enriched for this state whereas Capivasertib-rested organoids appeared more similar to parental (Fig. 4G).

Having established that AKT inhibition selects for cells in a basal state, we sought to test if cells in a basal state were intrinsically resistant to AKT inhibition. Drug-naïve cells were sorted on the basis of ITGA6 expression, which is high in basal cells, and subjected to AKT inhibition. Fig. S9E shows that drug-naïve ITGA6 high cells give rise to organoids that are less sensitive to AKT inhibition than ITGA6 low cells.

We investigated this observation more broadly. First, using DepMap, a publicly available dataset of cancer dependencies, we found basal-like breast cancer lines tended to be resistant to AKTi compared to luminal lines (Fig. S10A). In the presence of Capivasertib, resistant organoids studied here had slightly elevated levels of phosphoS6 compared to parental organoids (Fig. S10B) suggesting that PI3K pathway signalling may be altered in basal lines. Comparison of 25 luminal breast cancer cell lines and 8 HER2-enriched lines with 22 basal-like ones revealed that although PI3K inhibition reduces AKT activity in all three cell types, it has only a modest effect on S6 phosphorylation in basal-like lines (Fig. 4H & Fig. S10C). These data demonstrate that reduced coupling between AKT activity and S6 phosphorylation and growth control is a recurrent feature of breast basal-like cancer cells

### ΔNp63 is sufficient to confer resistance to AKT inhibition

We sought to identify regulators driving the transition into the basal drug resistant state. Analysis of genes enriched in Capivasertib-resistant basal organoids revealed a strong overlap with TP63 target genes (*32*). The transcription factor ΔNp63α regulates the basal cell lineage in the mammary gland (*33*). Overexpression of ΔNp63α, promoted elevated expression of basal cell markers ITGA6 and KRT14 and reduced levels of EPCAM, whereas the DNA-binding domain mutant (R304W) did not (Fig. S11A, B, C & D). Strikingly, ΔNp63α overexpression also rendered organoids insensitive to Capivasertib, whereas organoids expressing the R304W mutant were not (Fig. 4I). Thus, the pre-existing drug resistant state arises as a result of the endogenous program that determines basal cell identity.

### A basal-like cell state is enriched in Capivasertib-treated tumours

To directly test whether a Capivasertib-resistant state might make up a significant portion of a tumour we derived a 106-gene signature made up of ΔNp63α targets that were upregulated in Capivasertib-resistant organoids, which we name CaRe63. We interrogated our scRNA sequencing dataset of LARRY-labelled tumours and identified a subpopulation of cells that mapped to Cluster II and another that mapped to Cluster III, which was characterised by KRT14 expression (Fig. 5A and B). Gene set enrichment analysis of Cluster III markers identified basal-like breast cancer gene sets which were also identified in Capivasertib-resistant clusters *in vitro* (Fig. 5C). CaRe63 genes did not readily map onto any single GM reported by Wu et al. suggesting that they mark distinct cell states (Fig. S12A). Despite some similarity to a persister cell signature (Fig. S12B) (*34*), CaRe63 does not predict response to chemotherapy suggesting that the biology underpinning resistance to AKT inhibition is distinct from general chemoresistance (Fig. S12C).

**Fig. 5.**
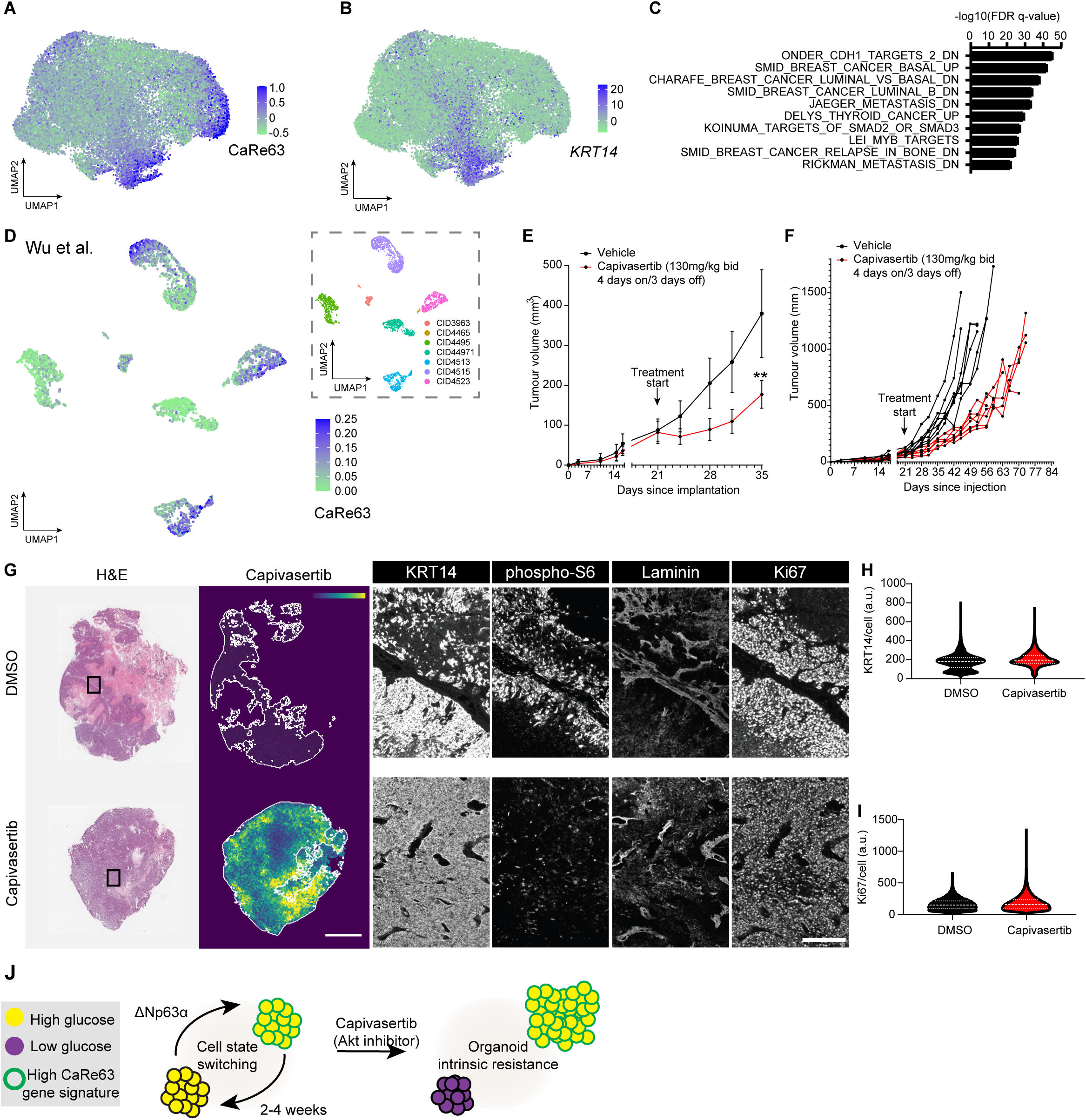
Capivasertib enriches for basal-like cells in TNBC tumours. **(A & B)** UMAP representation of GCRC1915 patient-derived xenograft single-cell RNA sequencing datasets with colour representing **(A)** upregulation of a gene signature defined by Capivasertib resistance and p63 target genes (CaRe63) and **(B)** *KRT14* **(C)** Barplot depicting gene set enrichment signature p-values from Cluster III marker genes. **(D)** UMAP representation of a human TNBC single-cell RNA sequencing dataset from Wu et al. with colour representing upregulation of a gene signature defined by Capivasertib resistance and p63 target genes (CaRe63) and (inset) patient identifier. **(E & F)** Tumour growth curves of GCRC1915 PDXOX derived tumours (**E** mean **F** individual) treated with either vehicle or Capivasertib (n=7 mice per group) ** p < 0.05. (Tukey’s multiple comparison test) **(G)** Multiplexed imaging micrographs of representative serial sections of vehicle or Capivasertib-treated tumours. **(H & I)** Violin plots of cellular staining intensity in DMSO and Capivasertib-treated tumours for **(H)** KRT14 and **(I)** Ki67 **(J)** Graphical model of cell state transitions and the impact of Akt inhibition on each of the cell states.

Interrogation of the cancer cells in TNBC patients described by Wu et al. revealed that some patients were uniformly low for CaRe63 (CID4495 & CID44971), while others exhibited intra-tumour heterogeneity similar to that observed in our experiments (CID4513, CID4515 & CID4523) (Fig. 5D). Variation in the proportions of CaRe63^+^ cells present in human tumours could be observed in 2 other independent patient cohorts (Fig. S12D & E) (*35, 36*). Based on our analysis we hypothesise that tumours lacking CaRe63 high cells would benefit from AKT inhibition, while those with a pre-existing population would rapidly become resistant.

Since GCRC1915 contains CaRe63 high cells, we predicted that this model would quickly become resistant to Capivasertib. Fig. 5E and F show that while Capivasertib significantly impaired tumour growth, this was only temporary and the tumour endpoint was only delayed by 3 weeks compared to control tumours. Using a combination of serial sectioning, mass spectrometry imaging and multiplex immunostaining we generated maps of Capivasertib and metabolite distribution, along with KRT14, phospho-S6, Laminin and Ki67 (Fig. 5G & Fig. S13A). Concordant with scRNA sequencing analysis, control tumours contained a mixture of KRT14 positive and negative cells (Fig. 5G). KRT14, as well as phospho-S6, positive cells were typically located near laminin, which denoted blood vessels and the edge of tumour cell nests. Both KRT14 positive and negative cells were positive for phosphoS6 and Ki67 staining indicating that they are both responsible for continued tumour growth. Capivasertib-treated tumours exhibited were dominated by KRT14 positive cells (Figure 5H). Mass spectrometry imaging confirmed that Capivasertib could be detected throughout the treated tumours and the increase in glucose levels indicated on target perturbation of physiological insulin signaling. Thus, inadequate drug delivery can be excluded as a reason for tumour growth. While we observed a decrease in phospho-S6 levels following AKT inhibition, Ki67 positivity was maintained (Fig. 5H), suggesting sufficient pathway activity for tumour growth. Similar to our *in vitro* analysis, metabolite profiling was similar between control and Capivasertib-resistant tumours (Fig. S13B & D). Interestingly, KRT14 positive cells were no longer confined to regions surrounding laminin staining but were uniformly distributed. These data establish that within 2-3 weeks Capivasertib treatment leads to the dominance of more basal KRT14 positive breast cancer cells.

Together, these analyses demonstrate that both *in vitro* and *in vivo* AKT inhibitor resistance is linked to the inherent biology of the breast, with basal cells exhibiting reduced coupling of AKT to S6 and regulation of glycolysis (Figure 5J).

## Discussion

The high frequency of mutations in *PI3K* isoforms, *PTEN*, and *AKT* make the pathway an attractive target for therapeutic intervention, however, Phase III clinical trials demonstrated overall efficacy of Capivasertib in ER+ disease, but not TNBC. To address this challenge, we implemented complementary approaches that overcome technological limitations that restrict lines of inquiry relating to biological variability in therapeutic responses. Using a 3D culture system of human TNBC organoids, LBETROSS and dOPM we have uncovered a non-genetic resistant cell state driven by ΔNp63α that is refractory to AKT.

The observed basal-like resistant state exhibited poor coupling between glucose metabolism and AKT activity. This may reflect differing requirements for extracellular cues in secretory luminal cells or contractile basal cells. Supporting this hypothesis, different metabolic sensitivities have been described for normal breast epithelial cell subtypes (*37*). In our model, this was recapitulated by the observed inter-organoid heterogeneity and the ability of ΔNp63 to promote AKT inhibitor resistance.

Using CaRe63 to define the cell state we believe that it is likely to pre-exist in many TNBC cases (*38, 39*) and, based on our analysis, we propose that patients CID4495 & CID44971 would benefit from Capivasertib, while others in the Wu et al. cohort would not. In tumours with CaRe63 high cells, it would be appealing to combine AKT inhibition with strategies that convert basal cell states to luminal. EZ2 inhibition in combination with AKT inhibition can trigger a cell death program in luminal cells with similarities to involution (*27, 34*); however, we did not observe synergy suggesting that it may not be sufficient to drive cells out of a basal state. We propose that future studies to identify therapeutic strategies to drive basal to luminal transitions will be valuable. These could overcome the resistance to AKT inhibition that we show results from inherently different coupling of AKT to cell growth between mammary cell states.

## Acknowledgements

The authors would like to acknowledge all members of the Sahai laboratory for their insightful comments. **C.D.H.R.** thanks the Cell Sciences, Flow Cytometry, Advanced Light Microscopy, Genomics, Bioinformatics and Biostatistics, Experimental Histopathology, High Throughput Screening, Human Biology Facility and Biological Research facilities from the Francis Crick Institute for their assistance and training. **C.D.H.R.** is a recipient of a Bourse postdoctorale from the Fonds de recherche du Québec – Santé (https://doi.org/10.69777/273104), as well as an EACR-AstraZeneca Postdoctoral Fellowship. **C.D.H.R.** is supported by funding from the AstraZeneca-Crick Research Alliance. **G.L.M.B.** is supported by an EMBO postdoctoral fellowship (ALTF-792-2021). **S.J.** acknowledges funding support from the BBSRC TRDF Grant (BB/T011572/1), Chris Banton Fund of The Francis Crick Institute, London and is the recipient of the LUSH Prize in the young researcher category. **S.J.** thanks MedTech Super Connector (MTSC) fellowship and BBSRC ICURe Explore Programme for funding support towards commercialization of innovation. **J.B.** is supported by the European Research Council under European Union (EU) Horizon 2020 research and innovation program grant 742138 and by the Wellcome Trust (220379/D/20/Z) **C.D.** is supported by EPSRC (EP/T003103/1), CRUK Accelerator (C10441/A29368) and Imperial EPSRC Impact Acceleration Account (EP/R511547/1) **E.S.** is supported by the Francis Crick Institute, which receives its core funding from Cancer Research UK (CC2040), the UK Medical Research Council (CC2040), and the Wellcome Trust (CC2040) and the European Research Council (ERC Advanced Grant CAN_ORGANISE, Grant agreement number 101019366). The breast cancer patient-derived xenograft organoids PDXO GCRC1915 was obtained from the breast tissue and data bank (Park lab, McGill University) (PMID: 32546838), supported by the Réseau de Recherche sur le Cancer of the Fonds de Recherche du Québec-Santé and the Québec Breast Cancer Foundation, and certified by the Canadian Tumor Repository Network (CTRNet). The work was supported by a CRUK Accelerator award (MACH3CANCER, A29368) and the dOPM instrument was initially developed under a project funded by the EPSRC (EP/T003103/1).

## Supplementary Materials

### Materials and Methods

#### Organoid lines, plasmids and compounds

GCRC1915 PDXO parental organoid lines were generated from the GCRC1915Tc PDX (Savage et al.). Organoids were routinely cultured in 12-well dishes pre-coated with a 150 µL layer of Cultrex RGF Basement Membrane Extract, type 2, PathClear®. Organoids were maintained (37°C, 5% CO_2_, humidified atmosphere) in culture media consisting of Advanced DMEM/F12 (Cat# 12634-010, Invitrogen), 1X GlutaMAX (Cat# 35050-061, Gibco), 10mM HEPES (Cat# 5630-080, Gibco), 1% Penicillin-Streptomycin (Cat# 15140-122, Gibco), 1X B-27 supplement (Cat# 17504044, Life tech), 0.1X mNoggin Conditioned Media (5X Concentrate obtained from Cell Services Platform at the Francis Crick Institute), 0.1X R-spondin-1 conditioned media (5X Concentrate obtained from Cell Services Platform at the Francis Crick Institute), 5 ng/mL human recombinant EGF (Cat# 78006.1, StemCell), 5 ng/mL human recombinant FGF-7 (KGF) (Cat# 78046, StemCell), 20 ng/mL human recombinant FGF-10 (KGF) (Cat# 78037.1, StemCell), 5 nM Neuregulin 1 (Cat# 78071.1, StemCell), 500 nM A83-01 (Cat# 2939/10, Tocris), 500 nM SB202190 (Cat# S7067, Sigma), 1.25 mM N-acetylcysteine (Cat# A9165, Sigma), 5 mM Nicotinamide (Cat# N0636, Sigma), 10 µM Y-27632 (Cat# 1254/1, Biotechne) and 5% Cultrex RGF Basement Membrane Extract, type 2, PathClear® (Cat# 3533-010-02, R&D Systems). Media was changed every 3 or 4 days and organoids were passaged once per week. For passaging, organoids were first treated with 1mg/mL collagenase/dispase diluted in resuspension media (DMEM/F12 (Cat# 11320033, Gibco) with 1% FBS and 1% Penicillin-Streptomycin (Cat# 15140-122, Gibco)) for 90 minutes. Then, organoids were resuspended in accutase for 10 minutes at room temperature, followed by 10 minutes at 37°C to obtain a single-cell suspension. Cells were pelleted, resuspended in resuspension media, counted and 1-2 × 10^5^ were plated in growth media. All organoid lines were authenticated and tested for mycoplasma contamination by the Cell Services Platform at the Francis Crick Institute. Clonal organoid lines were generated by single-cell sorting into 96-well v-bottom Cell-repellent plates (Cat#61970, Greiner Bio-One GmbH) followed by passaging after 2 weeks and expansion. Resistant organoid lines were generated by culturing organoids in the presence of 1 µM Capivasertib (Cat# HY-15431, Insight Biotechnology) for at least 1 month. The corresponding rested line was cultured without Capivasertib for 1-2 months.

##### Compound Table

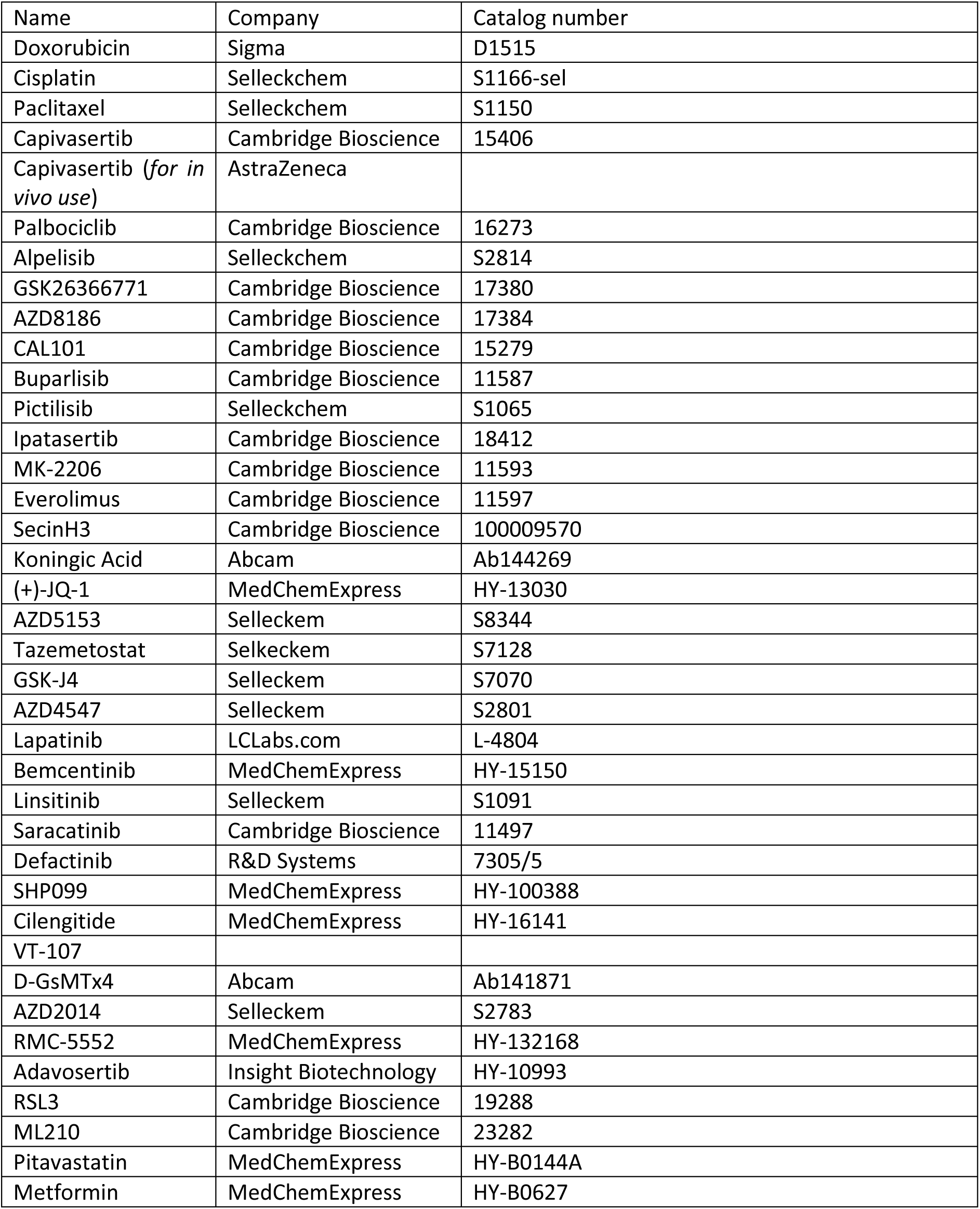
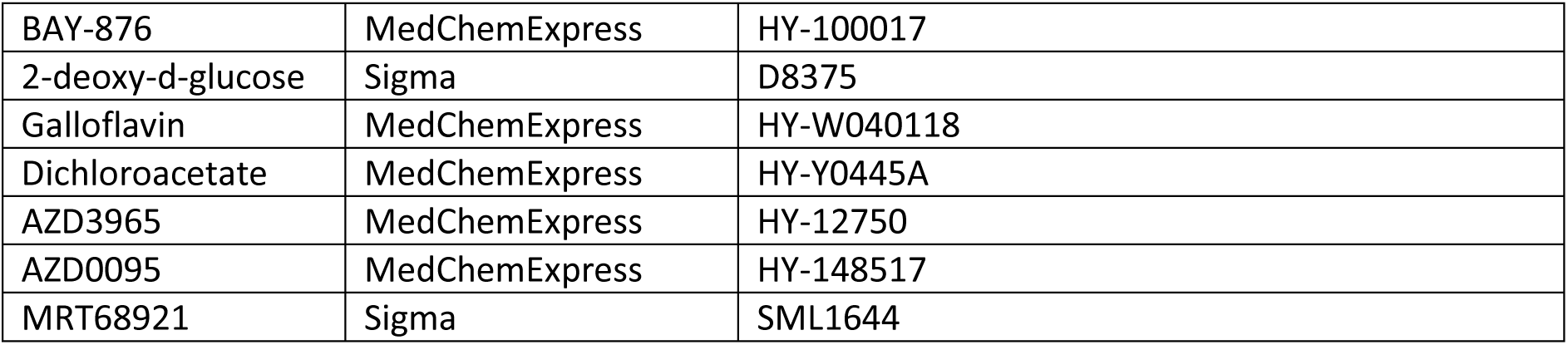

| Name | Company | Catalog number |
| --- | --- | --- |
| Doxorubicin | Sigma | D1515 |
| Cisplatin | Selleckchem | S1166-sel |
| Paclitaxel | Selleckchem | S1150 |
| Capivasertib | Cambridge Bioscience | 15406 |
| Capivasertib (*for in vivo use*) | AstraZeneca |  |
| Palbociclib | Cambridge Bioscience | 16273 |
| Alpelisib | Selleckchem | S2814 |
| GSK26366771 | Cambridge Bioscience | 17380 |
| AZD8186 | Cambridge Bioscience | 17384 |
| CAL101 | Cambridge Bioscience | 15279 |
| Buparlisib | Cambridge Bioscience | 11587 |
| Pictilisib | Selleckchem | S1065 |
| Ipatasertib | Cambridge Bioscience | 18412 |
| MK-2206 | Cambridge Bioscience | 11593 |
| Everolimus | Cambridge Bioscience | 11597 |
| SecinH3 | Cambridge Bioscience | 100009570 |
| Koningic Acid | Abcam | Ab144269 |
| (+)-JQ-1 | MedChemExpress | HY-13030 |
| AZD5153 | Selleckem | S8344 |
| Tazemetostat | Selleckem | S7128 |
| GSK-J4 | Selleckem | S7070 |
| AZD4547 | Selleckem | S2801 |
| Lapatinib | LCLabs.com | L-4804 |
| Bemcentinib | MedChemExpress | HY-15150 |
| Linsitinib | Selleckem | S1091 |
| Saracatinib | Cambridge Bioscience | 11497 |
| Defactinib | R&D Systems | 7305/5 |
| SHP099 | MedChemExpress | HY-100388 |
| Cilengitide | MedChemExpress | HY-16141 |
| VT-107 |  |  |
| D-GsMTx4 | Abcam | Ab141871 |
| AZD2014 | Selleckem | S2783 |
| RMC-5552 | MedChemExpress | HY-132168 |
| Adavosertib | Insight Biotechnology | HY-10993 |
| RSL3 | Cambridge Bioscience | 19288 |
| ML210 | Cambridge Bioscience | 23282 |
| Pitavastatin | MedChemExpress | HY-B0144A |
| Metformin | MedChemExpress | HY-B0627 |
| BAY-876 | MedChemExpress | HY-100017 |
| 2-deoxy-d-glucose | Sigma | D8375 |
| Galloflavin | MedChemExpress | HY-W040118 |
| Dichloroacetate | MedChemExpress | HY-Y0445A |
| AZD3965 | MedChemExpress | HY-12750 |
| AZD0095 | MedChemExpress | HY-148517 |
| MRT68921 | Sigma | SML1644 |

##### Primers

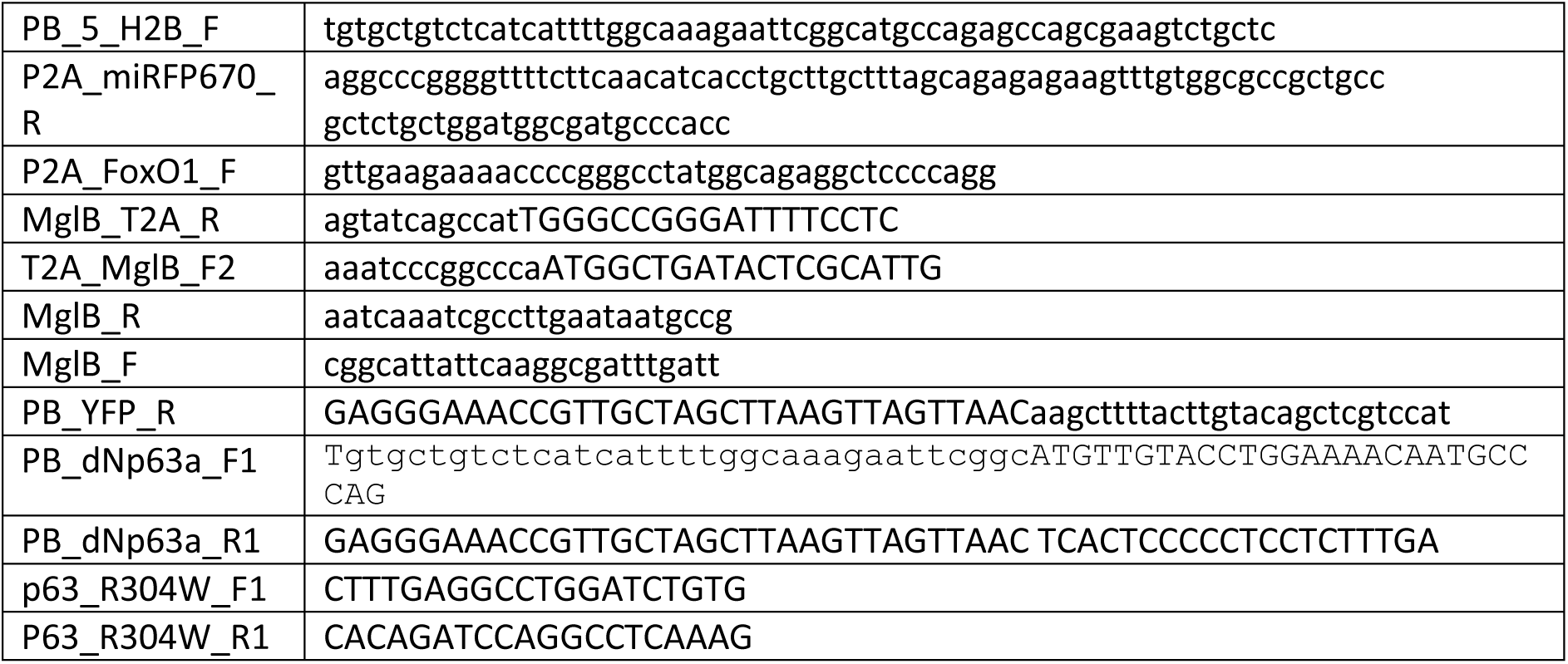

Stable organoid lines were generated using the piggybac transposon system. A 1:1 mix of relevant and pBase plasmids (4 µg total DNA) and 12 µL Lipo2000 were incubated for 5 minutes and then added to 0.5 × 10^5^ resuspended cells. This mix was incubated (37°C, 5% CO_2_, humidified atmosphere) in an ultra-low adherent plate (Cat#10154431, VWR) for 1-4 hours and then replated under standard conditions. After 4 days, media was changed and organoids were selected and continuously cultured with 5 µg/mL blasticidin.

Constructs H2B-miRFP670nano (Cat# 127438, Addgene), pSBbi-FoxO1_1R_10A_3D (Cat# 106278, Addgene) and ΔNp63α were kind gifts from Vladislav Verkhusha, Laura Heiser and Karen Vousden respectively. eCFP-MglB-Citrine was described previously. The polycistronic (H2B-miRFP670nano-P2A-FoxO1_1R_10A_3D-T2A-eCFP-MglB-Citrine), mEmerald, ΔNp63α and ΔNp63α (R304W) expressing vectors was generated using PCR amplified fragments and cloned into EcoR1-HF and Hpa1 digested piggybac transposon vector by Gibson Assembly (NEB). All constructs were sequence verified before use.

#### *In vivo* experiments

The Francis Crick Institute’s Animal Welfare and Ethical Review Body and UK Home Office authority provided by Project License 0736231 approved all animal model procedures. Procedures described in this study were compliant with relevant ethical regulations regarding animal research. For drug studies, Capivasertib was formulated once weekly as a solution in 10% DMSO, 25% Kleptose and dosed twice daily (BID) 8 hours apart by oral gavage 0.1 mL/10g of the animal using a schedule of 4 days dosing, 3 days not dosing at 130 mg/kg. Control animals were dosed with vehicle equivalent to compound dosing groups.

#### Organoid Lineage tracing

##### LARRY library amplification

LARRY Barcode Version 1 library was a gift from Fernando Camargo (Addgene #140024). 20 ng of the LARRY plasmid library was transformed into 12 vials (25 µL each) of Stbl4 ElectroMAX cells (ThermoFisher, 11635018). Following 1-hour recovery at 37°C in SOC medium, each bacterial tube was plated onto two 20 × 20 cm agar plates. The next day, cells were rinsed 3 times from plates into a total of 1.5 L of LB, cultured for 2.5 hours at 37°C, and purified using the NucleoBond Xtra Maxi EF (Macherey Nagel, 740424.50). The final DNA yield was 1–1.5 mg.

##### Lentivirus production

Lentiviral particles were produced using a third-generation lentivirus packaging system: pLP1 (Invitrogen, A43237), pLP2 (Invitrogen, A43237) and pVSV-G (System Biosciences). LARRY library and packaging plasmids were transfected into HEK293T, cells using Lipofectamine 2000 (Invitrogen, 11668500). HEK 293T cells were grown in Dulbecco’s modified Eagle’s medium (DMEM) (Sigma, D5796) supplemented with 10% foetal bovine serum (FBS) (Biosera, FB-1001/500) and 1% penicillin/streptomycin (Gibco, 15140-122) and transfected in Opti-MEM I (Gibco, 31985). Sixteen hours after transfection, media was replaced with Dulbecco’s modified Eagle’s medium (DMEM) (Sigma, D5796) supplemented with 10% fetal bovine serum (FBS) (Biosera, FB-1001/500) and 1% penicillin streptomycin (Gibco, 15140-122). 36 and 60 hours after transfection, media were collected, passed through a 0.45 μM filter (Millex-HV Millipore, SLHV033RS), and then ultracentrifuged at 22,000 rpm for 1.5 hours. Pellets were resuspended in sterile Dulbecco’s phosphate-buffered saline (DPBS) (Gibco, 14190) and stored at −80 °C.

##### Plasmid and lentivirus library diversity assessment

For plasmid library sequencing, primers pLARRYfw (TCGTCGGCAGCGTCAGATGTGTATAAGAGACAGaggaaaggacagtgggagtg) and pLARRYrv (GTCTCGTGGGCTCGGAGATGTGTATAAGAGACAGcaaagaccccaacgagaagc) were used to amplify the plasmid library. For lentiviral library sequencing, RNA was extracted from 40 µL of the viral preparation using the Virus NucleoSpin Kit (Macherey Nagel, 740983.50), eluted in 40 µL. 3 µg of RNA were retrotranscribed to cDNA with Superscript VILO (random primers, ThermoFisher 11754050). Amplification was performed with primers vLARRYfw (TCGTCGGCAGCGTCAGATGTGTATAAGAGACAGaatcctcccccttgctgtcc) and vLARRYrv (GTCTCGTGGGCTCGGAGATGTGTATAAGAGACAGaccgttgctaggagagaccata.

In both cases, mock qPCR was used to determine the minimum cycles to avoid saturation. PCR amplification was then conducted with Q5 High-Fidelity Polymerase (NEB, M0494) using the following conditions: 98°C for 30 s; [98°C for 10 s, 63°C for 30 s, 72°C for 20 s] x 7–15 cycles as per qPCR results; 72°C for 2 min. Libraries were size-selected (0.5–0.2X dual-sided) with Ampure XP beads (Beckman Coulter, A63881) and sequenced to a depth of ∼5 million reads (plasmid) or ∼10 million reads (virus).

Sequencing data were analysed using the genBaRcode R package (v1.2.5), yielding a library diversity of approximately 0.25 million barcodes for both libraries, consistent with the original library diversity reported on Addgene.

##### Organoid infection

3 × 10^4^ cells were aliquoted and incubated with 1.5 × 10^6^ LARRY virus particles in 200 µL culture media lacking Cultrex under culture conditions (37°C, 5% CO_2_, humidified atmosphere) for 1 hour. Cells were then split into 3 and plated in 12-well dishes pre-coated with a 150 µL layer of Cultrex RGF Basement Membrane Extract, type 2, PathClear®. After 3 days, media was removed and replaced with 1mL of indicated culture media and replaced in culture conditions. After 4 more days, approximately 30% of organoids were eGFP+ and were processed further.

##### Organoid enrichment using the VISIBLE system and tumour processing

The VISIBLE system comprises two independent units: a top unit, which features a robotic manipulation system with two interchangeable tool heads, and a bottom unit that is an automated fluorescence microscope. Briefly, the robotic manipulation consists of an XY gantry with tool-head and its motion is controlled by 2 stepper motors (Bipolar Hybrid Stepper Motor Nema 17) arranged in a core XY coordination. For motion control in Z axis, a customized in-house developed screw rail-based mechanism is employed with a stepper motor (Bipolar Hybrid Stepper Motor Nema 11). Mechanical limit switches are installed for homing the axes. Stepper motors and limit switches are controlled with firmware installed on Smothieboard V2. We have used open-source Pronterface software module for interactively controlling our robotic system. The bottom unit is a fluorescence inverted microscope procured from Zaber Technologies Inc with a 2.3-megapixel CMOS sensor (Hamamatsu ORCA-Spark C11440-36U). Images were acquired using a FITC filter (Semrock BrightLine full-multiband filter set, optimized for DAPI, FITC, & Texas Red) and the light source consists of 6 solid-state sources operating independently from Lumencor (SPECTRA X) with output spectral range between 360–780 nm. A 3 mm liquid light guide was used to connect the light source with the microscope chamber. Open-source micro-manager software was used to control and automate the microscope unit.

Using this system, eGFP+ organoids were picked, spun down, and resuspended in Matrigel (Corning Cat No. 354234) prior to injection into the 4^th^ inguinal mammary gland. 5 weeks after injection of eGFP+ enriched organoids, tumours were collected, dissociated and human tumour cells were enriched using a Miltenyi tumour dissociator according to manufacturer’s instructions. Briefly, tumour tissues were minced into 0.5-1.0 mm fragments with a sterile razor blade and this was transferred to a gentleMACS C tube (Order No. 130093-237). 4.7 mL DMEM was added along with enzymes A,R and H prior to running the 37C-h-TDK3 program for 30 minutes. Samples were then incubated on ice for 5 minutes and 3-4 mL of supernatant was passed through a 70 μm strainer. The remaining tumour pieces were broken apart using the m-impTumor-01-01 program for 1 minute and this solution was passed through the same 70 μm strainer. The strainer was washed once with PBS and collected cells were pelleted by centrifuging at 1200 rpm for 5 minutes at 4°C, washed once with PB in 80 μL PBS+0.5% BSA. Cell suspension was incubated with 20 μL Miltenyi Mouse Cell Depletion cocktail antibody for 15 minutes at 4°C. Volume was adjusted to 500 μL and passed through a Miltenyi LS column (Order No. 130-042-401). Flow through containing enriched human tumour cells were collected and used subsequently for single cell RNA sequencing.

##### Organoid sorting

eGFP+ organoids were sorted by first treating with collagen and dispase followed by resuspending in 30 mL FACS buffer (1% BSA in PBS w/v) and sorting using a complex object parametric analyser and sorter (COPAS) VISION 250 particle sorter. The gating strategy used to identify LARRY expressing organoids was based on the optical density (Extinction 561 peak height), size (time of flight) and eGFP fluorescence (488 nm laser excitation and detected using a 512/26 bandpass filter). A neutral density ND 2.0 filter was used to prevent extinction signal saturation and the gating strategy was confirmed based on in-flow brightfield images. Sorted eGFP+ organoids were centrifuged and digested with accutase to obtain a single-cell suspension.

##### Single cell RNAsequencing

Single cell concentration and viability was measured using acridine orange (AO) and propidium iodide (PI) and the Luna-FX7 Automatic Cell Counter. Approximately 120 000 cells per tumour or 900-7300 cells per LEBTROSS sample were loaded on Chromium Chip and partitioned in nanolitre scale droplets using the Chromium Controller and Chromium Next GEM Single Cell Reagents (10x Genomics CG000315 Chromium Single Cell 3’ Reagent Kits User Guide (v3.1 - Dual Index)). Within each droplet the cells were lysed, and the RNA was reverse transcribed. All of the resulting cDNA within a droplet shared the same cell barcode. Illumina compatible libraries were generated from the cDNA using Chromium Next GEM Single Cell library reagents in accordance with the manufacturer’s instructions. Final libraries are QC’d using the Agilent TapeStation and sequenced using the Illumina NovaSeq 6000. Sequencing read configuration: 28-10-10-90.

##### Data processing and analysis

The GFP-LARRY plasmid was added to the reference genome refdata-gex-GRCh38-2020-A and the GTF annotation of the plasmid transcripts was added to the same reference annotation file. Cellranger (version 7.0.1) mkref was ran to create a new reference for genome an annotation. Cellranger count was used to count the 10x libraries.

The LARRY barcode information was obtained and extracted from the 10X GEX library and from the libraries containing the specifically amplified LARRY barcodes. To extract the unmapped reads and the reads that map to the LARRY-GFP plasmid (mRNA) from the 10X/CellRanger output, we identify those reads using samtools (version 1.15.1) (*40*) and subsequently extract those reads from the 10X GEX fastq.gz files. The LARRY barcodes are extracted from the LARRY libraries by removing primer/adapter sequences with cutadapt (version 3.5) (*41*) followed a second round of cutadapt to extract the LARRY barcodes.

A whitelist of cell barcodes was obtained with UMI-tools (version 1.1.2) (*42*), specifically umi_tools extract, and after having ensured that each unique cell-barcode/UMI combination is associated with one LARRY barcode sequence, we grouped all the LARRY barcodes found within the same cell. For each cell barcode-LARRY barcode combination, we count the number of unique UMIs associated with each cell barcode-LARRY barcode combination. After having identified a set of Cell barcode-LARRY barcode - UMI counts for each of the samples, we next identify clonal groups for all samples.

The cellranger quantifications were imported into a Seurat (version 4.3.0) (*43*) object and doublets were calculated using scDblFinder (version 1.8.0)(*44*) and the LARRY barcode information was added to the Seurat objects. The R programming language was used (version 4.1.2)(R Development Core Team 2008). We investigated key QC parameters, and we remove cells with a low number of detected features or cells with a very high proportion of mitochondrial gene expression. We selected a lower bound filter for both the minimum number of reads per cell and the minimum number of detected features using 3 median absolute deviations (MADs). For the percentage of mitochondrial gene, we set upper bounds using 3 MADs.

Before investigating clonal groups across cluster (used as a proxy for cell state) we remove cells that are associated with more than one clonal group (i.e. LARRY barcode).

We use the “SCTransform” method (version 0.3.5)(*45*) for the normalisation and variance stabilisation which, in comparison to the standard Seurat approach, sctransform returns 3000 variable features by default, instead of 2000.

Clonal groups are identified across all samples based on the combined library prep information for each sample. We investigate the percentage of cells in which LARRY barcodes were detected, the number of unique clonal groups, the number of detected clonal groups in cells with detected LARRY barcode, the number of cells within each clonal group.

#### Analysis of CaRe63 in single cell RNA sequencing datasets

Analysis was performed using the Seurat package (version 4). Gene modules were obtained from Wu et al. and data downloaded from GEO GSE176078. Kim et al. data was obtained from SRA: SRP114962 and Biokey from EGA: EGAS00001004809. To assess the over-representation of the CaRe63 signature in various gene sets, a hypergeometric test was performed using the phyper function from the stats R package (v4.1.0). Let *n* be the number of genes in the CaRe63 signature, *m* the number of genes in the queried gene set, *i* the number of overlapping genes between CaRe63 and the queried set, and *U* the total number of genes in the universe. Cumulative probability of the right tail of the hypergeometric distribution is used for over-representation statistical significance and is calculated as follows.

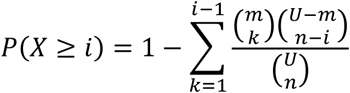

*P* values were adjusted for multiple testing using the Benjamini–Hochberg false discovery rate (FDR) method. Gene sets with an adjusted P < 0.05 and at least three overlapping genes were considered significantly enriched for CaRe63.

#### SCAN-B survival analysis

Bulk RNA-seq data from the Sweden Cancerome Analysis Network – Breast (SCAN-B) clinical cohort was retrieved (*46*). Survival analysis was conducted on a subset of chemotherapy-treated TNBC patients (n = 231). CaRe63 scores of patients were derived by calculating the first principal component (PC1) of the expression levels of CaRe63 genes. Patients were stratified into high and low CaRe63 groups based on the median of PC1 value.

Kaplan–Meier curves for overall survival (OS) over a 10-year period post-diagnosis were generated using the survdiff and coxph functions from the survival R package (v3.2.13). The coxph function fits a Cox proportional hazards regression model, and survdiff performs a log-rank test to evaluate the statistical significance of the survival differences between the CaRe63-low and CaRe63-high groups.

#### Incucyte imaging and organoid growth analysis

Organoids were generated by plating 1 × 10^4^ cells per well in a 48-well dish (Cat# 353078, FALCON) pre-coated with a 50 µL layer of Cultrex RGF Basement Membrane Extract, type 2, PathClear®. After 3 days, culture media was changed to culture media containing the indicated treatments. The plate was transferred to an Incucyte S3 live cell imaging instrument contained within a standard tissue culture incubator and allowed to equilibrate for 1-2 hours. Images were acquired with a 10X objective every 3 hours for 4 days to capture 33 time points. Within FIJI, images were made into stacks, stabilized and individual organoids were tracked using the TrackMate batcher plugin. Within this plugin, CellPose 2.0 was used to segment individual organoids and was used to identify tracks that lasted the entire length of the acquisition. Exponential growth curves were then fitted to each growth curve and a rate was determined for each organoid using custom Python code. Plots were then generated using the SuperPlotsOfData Shiny app or Graphpad.

#### Power calculation for modified Luria-Delbrück

The probability, or confidence, that none of the selected clones were resistors was given by:

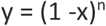

Where x is the probability of being a resistor and n is the number of clones selected. Solving for n, we found that from a population where 17% of cells are resistant, we would need to test at least 17 clonal populations to have a > 95% confidence that resistance is not a clonal feature.

#### Dual-view oblique plane microscopy imaging

##### Image acquisition

Organoids were generated by plating 1 × 10^4^ cells per well in a ibidi 24 well µ-plate (Cat# 82426, ibidi) pre-coated with a 83 µL layer of Cultrex RGF Basement Membrane Extract, type 2, PathClear®. After 3 days, culture media was changed to phenol-red free imaging media. Imaging media consisted of SILAC Advanced DMEM/F-12 Flex, supplemented with 17.5 mM D-Glucose, 0.699 mM L-Arginine hydrochloride and 0.499 mM L-Lysine hydrochloride, along with additional supplements present in culture media. Organoids were imaged on a custom-built dual-view oblique plane microscopy (dOPM) system as previously described(*47, 48*). For each field of view, two dOPM views were acquired at ±35°, with z-planes spaced 1 μm apart and spanning a 150 μm depth (perpendicular to the imaging plane). Images were collected at 512 × 512 resolution with a pixel size of 0.35 μm in the sample space. The light-sheet width was approximately 3 μm FWHM at the waist. Two dOPM views were deskewed and fused using the Multiview Fusion plugin in FIJI (*49*), with affine registration derived from 3 μm UltraRainbow fluorescent beads (Spherotech, URFP-30-2) embedded in 2–4% agarose. Final volumes were converted to OME-TIFF format for further analysis. Two fused views were resliced and binned ×2 to produce isotropic 0.69 × 0.69 × 0.69 μm³ voxels.

H2b, Akt activity and glucose biosensor were imaged using five fluorescence channels: eCFP (445 nm excitation, 483/32 nm emission), sensitized Citrine-FRET (445 nm/525/45 nm), direct Citrine (515 nm/542/20 nm), mRuby2 (561 nm/609/54 nm), and miRFP670nano (642 nm/697/58 nm). Each organoid was imaged immediately before treatment, 24, and 96 hours after drug treatment. The detection camera rolling shutter exposure time was set to 10 ms for all channels. All laser power levels were set to 100% within Omicron laser control software and NIS-elements acquisition software apart from 561 nm excitation which was set to 20% of maximum power. For each excitation wavelength, average power in the back focal plane was measured to be: 445 nm @ 2.4 mW, 515 nm @ 3.2 mW, 561 nm @ 0.8 mW and 642 nm @ 5.6 mW. All lasers were blanked so as to only be active during the camera exposure time.

Fixed organoids were processed according to the FLASH immunofluorescence protocol described for organoids (*50*). Briefly, organoids were fixed with a 4% paraformaldehyde solution, washed 3 times with PBS, FLASH reagent 2 was used for permeabilization, followed by washing with PBT, blocking, and overnight primary staining at 4 °C (1:1000, Krt14 Clone LL002 Cell Services Platform at the Francis Crick Institute). Organoids were washed with PBS, and stained with secondary (1:2000 Goat anti-rabbit 555 (Cat# A21428 Invitrogen)) for 1 hour at room temperature. Nuclei were counterstained with 0.5 µmL siR-DNA in PBST for 5 minutes at room temperature, followed by washing with PBS. Fluorescence imaging used 488 nm (Krt14, 525/45 nm) and 642 nm (nuclear, 697/58 nm) channels. Imaging used 20 ms exposure and 100% laser power for all channels.

##### Image processing

For each fluorescence channel, the background was removed using a threshold set at 1.4 times the background signal level determined from the mode of each image volume. For the glucose/AKT data, the nuclear channel was log-transformed.

Nuclear segmentation was performed using a nonlinear multiscale top-hat enhancement transform following (*51, 52*). This was performed by first smoothing with 3D Gaussian filters at three spatial scales of σ₁ ≈ 1.4 μm, σ₂ ≈ 2.8 μm, σ₃ ≈ 5.6 μm (corresponding to kernel sizes of 2, 4, and 8 voxels at 0.7 μm/voxel), resulting in images U_1_, U_2_ and U_3_ respectively. The enhancement used two paired scale comparisons: the normalized difference between U₁ and U₂, i.e. (U₁−U₂)/U₂, and between U₂ and U₃, i.e. (U₂−U₃)/U₃. These responses were combined linearly with a fixed mixing weight (a₁ = 0.7) to prioritize fine-scale structure. A global threshold was then applied to generate a binary mask. Threshold values were fixed per experiment (FUCCI, glucose/AKT, keratin) and tuned manually by inspecting representative volumes to optimize segmentation quality by eye.

Small holes were filled and thin structures removed using morphological operations. Objects smaller than ∼3.5 μm in diameter (i.e. <125 voxels) were excluded. A 3D watershed refinement step was applied to separate touching nuclei, and objects at the volume border or outside the main organoid region were discarded.

In experiments with cytoplasmic biosensors (FRET, AKT, keratin), cytoplasmic “collar” regions were defined for each nucleus by performing two successive 1 μm 3D dilations of the nuclear mask. The collar was then extracted as the shell-like region between the second dilation and the original nucleus, excluding any areas overlapping nuclear or collar regions from neighbouring cells. Biosensor intensities were then quantified separately within nuclear and cytoplasmic compartments.

##### Feature Extraction

For each segmented nucleus and corresponding collar, summary statistics were computed for each fluorescence channel, including the mean, standard deviation, skewness, kurtosis, and 25th/50th/75th percentiles. FRET ratio (Citrine/CFP) was calculated as the ratio of corrected Citrine intensity to corrected CFP intensity. AKT activity was defined as:

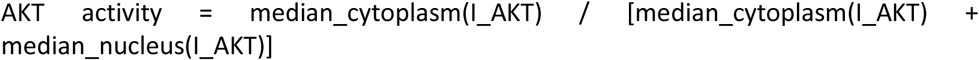

*where I_AKT* refers to the background-corrected AKT intensity, and “median_cytoplasm” and “median_nucleus” denote the median intensity values within cytoplasmic collars and nuclei, respectively.

Voxel-wise biosensor overlays were generated by assigning per-object median values back into the segmented binary masks. These overlays were saved as five-channel OME-TIFF hyperstacks containing raw intensities, biosensor activity maps, and segmentation masks.

##### Proliferation Rate Estimation

To determine the proliferation rate of each organoid, the number of segmented nuclei per organoid was compared between timepoints. The per-organoid proliferation rate was calculated as:

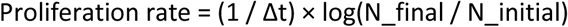

where Δt is the time interval in hours, N_initial is the number of nuclei at the starting time point, and N_final is the number at the later time point.

#### Bulk RNA sequencing

Organoids were generated by plating 3.5 × 10^4^ cells per well of a 12-well plate. After 3 days, media was removed, cells lysed and mRNA isolated according to manufacturer’s instructions, including on column Dnase digest (QIAGEN Rneasy mini kit).

Sequencing libraries were prepared using the KAPA mRNA HyperPrep Kit (Roche, kit code KK8581) according to manufacturer’s instructions. Briefly, samples were normalised to 50 ng of RNA in a final volume of 50 µl. PolyA-tailed RNA was captured with 50 ul of capture beads at 65 °C for 2 min and 20°C for 5 min. Beads were washed to remove other RNA species and RNA was eluted from the beads in 50 ul of RNase free water by incubating at 70 °C for 2 min and 20 °C for 5 min. A second capture of PolyA-tailed RNA was performed by adding 50 µl of bead binding buffer and the sample incubated for 5min at 20 °C. Beads were washed to remove other RNA species and eluted in 20 µl of Fragment, Prime and Elute Buffer. The fragmentation reaction was run for 6 min at 94 °C for a library insert size of 200-300 bp. Fragmented RNA underwent first and second strand cDNA synthesis according to manufacturer instructions. The KAPA Unique Dual-Indexed Adapters Kit (15μM) (Roche-KK8727) stock was diluted to 1.5 µM as recommended by the manufacturer for a total RNA input of 50 ng. The adaptor ligation was carried out by adding 5 µl of diluted adaptor and 45 µl of ligation mix (50 µl of ligation buffer + 10 µl of DNA ligase) to 60 µl of cDNA. The ligation reaction was run for 15 mins at 20°C. To remove short fragments such as adapter dimers, two SPRISelect bead clean-ups were done (0.63x SPRI and 0.7x SPRI). To amplify the library, 25 µl of Kapa HiFi HotStart PCR master mix plus 5 µl of Library Amplification Primer mix was added to 20 µl cDNA and 14 PCR cycles were performed as recommended by the manufacturer for a total RNA input of 50 ng. Amplified libraries were purified via a SPRISelect 1x bead cleanup.

The quality and fragment size distributions of the purified libraries was assessed by a 4200 TapeStation Instrument (Agilent Technologies). An average of 25 million paired end reads per library (PE100) were generated on a NovaSeq6000 (Illumina). All samples were processed using the nf-core RNAseq pipeline (v.3.10.1) operating on Nextflow (v.22.10.3) (68, 69). Settings for individual tools were left as standard for this workflow version unless otherwise specified. Alignment to GRCm38 (release 95) was performed using STAR (v2.7.10a) (64), followed by quantification using RSEM (v1.3.1) (63). Results were further processed in R (version 4.2.0) (67). Outputs of the RNAseq pipeline were assessed for quality using the metrics provided by the pipeline’s inbuilt quality control packages, PCA (package stats v4.2.0), and correlation analyses between samples (package stats v4.2.0). Differential expression analysis was performed with DESeq2 (v1.38.3) (66), using a model accounting for differences in replicate and treatment (∼ replicate + treatment). The significance threshold for the identification of differentially expressed genes was set as an adjusted P value ≤0.05, where P values were adjusted for false discovery rates according to the Independent Hypothesis Weighting method (package IHW, v1.26.0) (70, 71). Bulk RNAseq data have been deposited in the Gene Expression Omnibus, accession codes: XXXX and XXXX

#### Metabolomics

##### Sample preparation

Organoids were generated by plating 150 000 cells per well into an ultra-low attachment 6-well plate (Corning ref. 3471) in organoid culture media described above. After 3 days of culture, organoids were collected, spun down (1300 rpm for 4 mins), rinsed with SILAC Advanced DMEM/F12 without glucose and resuspended in 2 mL Breast Cancer Organoid Media without glucose that was supplemented with 17.5 mM ^13^C-Glucose, 91.25 mg/L L-lysine and 147.5 mg/L L-Arginine. Organoids were then replated on ultra-low attachment 6-well plates and incubated at 37*°C and 5% CO_2_*. After 6 hours, organoids were collected, rinsed with 10 mL chilled PBS and put on ice. These were centrifuged for 3 minutes at 600 g in cooled centrifuge and the supernatant along with as much of the matrigel layer as possible was aspirated without touching the cell pellet. 800 uL cold PBS was added and organoids were transferred to a chilled 2 mL Eppendorf tube. Centrifuge for 3 minutes at 600 g in cooled centrifuge. Pipette off the supernatant and add 900uL MeOH containing 1 nmol scyllo-inositol. Freeze on dry ice and samples were stored -80*°C*. To extract metabolites, add 300 uL CHCL3 followed by 300 uL H20 to each sample. Vortex briefly but vigorously and pulse sonicate in a sonicating water bath in a cold room. Incubate the samples for 1 hour (8 minutes ON, 7 minutes OFF for four cycles). Then, centrifuge for 10 minutes at maximum speed (16 100g). Transfer supernatant to 2 mL eppendorf and store at -80. The remain cellular debris was used for DNA extraction using the PureLink Genomic DNA kit (ThermoFisher); quantify total DNA content coming from each sample using a NanoDrop. Prior to GC-MS, dry metabolomic samples in speed vac. Each extract will be partitioned using MeOH/Water/CHCl3 (150:150:50), vortexed briefly, centrifuge at 4*°C* for 10 min.

##### Gas Chromatography-Mass Spectrometry (GC-MS)

Data acquisition was performed using an Agilent 7890B-5977A GC-MSD in EI mode after derivatization of twice methanol-washed dried extracts by addition of 20 μL methoxyamine hydrochloride (20 mg/mL in pyridine (both Sigma), RT, >16 hrs) and 20 μL BSTFA + 1% TMCS (Sigma, RT, >1 hr), for polar metabolites. GC-MS parameters were as follows: carrier gas, helium; flow rate, 0.9 mL/min; column, DB-5MS+DG (30 m + 10 m × 0.25 mm, Agilent); inlet, 270°C; temperature gradient, 70°C (2 min), ramp to 295°C (12.5°C/min), ramp to 320°C (25°C/min, 3 min hold). Scan range was m/z 50-550 (polar). Data was acquired using MassHunter software (version B.07.02.1938). Data analysis was performed using MANIC software, an in house-developed adaptation of the GAVIN package (PMID: 21575589). Metabolites were identified and quantified by comparison to authentic standards, and label incorporation estimated as the percentage of the metabolite pool containing one or more 13C atoms after correction for natural abundance.

#### Flow cytometry

Multiparametric flow cytometry was used for analysis of organoid cells. Organoids were dissociated and Cells were stained with a blue fixable Live/Dead dye (Invitrogen; Thermo Fisher Scientific) for the exclusion of dead cells before incubation with saturating concentrations of surface monoclonal antibodies (mAbs ADD details of the antibodies) diluted in 50% Brilliant violet buffer (BD Biosciences) and 50% PBS for 30 min at 4°C. Cells were fixed and permeabilized for further functional assessment with Cytofix/Cytoperm (BD Biosciences) according to the manufacturer’s instructions. All samples were acquired in 1x PBS using a Cytek Aurora and analyzed using FlowJo v10.10.0 (Tree Star). Doublets and dead cells were excluded from the analysis.

#### Comparison of PI3K inhibition in luminal and basal cell lines

Single-cell mass cytometry time of flight data from Tognetti et al.(*53*) was loaded from the corresponding Synapse repository (Project SynID: syn20366914) using the Python *synapseclient* API. Loaded files included: all those in the ‘single_cell_phospho’ (syn20613592) folder apart from those in the ‘subchallenge_4’ subfolder; *sc1gold* (syn20631271), *sc2gold* (syn20631273) and *sc4gold* (syn20631275) in the ‘goldstandard’ subfolder. The *FileID_table* file (syn20631269) was also loaded to obtain information on time course acquisition date and replicate. All files apart from *sc4gold* were processed together, computing the pseudo-bulk response by averaging across all single cells belonging to the same cell line, under the same treatment condition, in the same timepoint and replicate and with the same file ID. For cell lines in subchallenges 1 and 2, missing entries – removed for the purposes of the 2019 DREAM Challenge, see Gabor et al.(*54*) – were dropped and replaced with the full entries from the corresponding gold standards (*sc1gold* and *sc2gold*, respectively). As data for cell-lines in subchallenge 4 (*sc4gold*) is already provided in the form of population-level response, it was directly pivoted into the same format and concatenated with the processed data from above.

The resulting data was subset to the markers of interest (phospho-AKT, both Thr308 and Ser473, and phospho-S6) and treatments of interest (stimulation with EGF in the absence or presence of the pan-PI3K inhibitor pictlisib). Subtype annotations were obtained from Marcotte et al. (*55*) (https://github.com/neellab/bfg/tree/gh-pages/data/annotations) using the so-called intrinsic subtype unless otherwise specified (‘subtype_intrinsic’). Cell lines DU4475, HCC2157 and MDA-kb2 – for which annotations were missing – were annotated as Triple Negative Breast Cancer (TNBC) using Cellosaurus (*56*). Cell lines HDQ-P1 and MA-CLS-2, for which conflicting annotations were found, were re-annotated as TNBC following the subtype information in Tognetti et al. and other subtype categorisations in Marcotte et al., namely three-receptor status (‘subtype_three_receptor’) and Neve subtype (‘subtype_neve’). Subtype annotations were further aggregated, grouping Luminal A, Luminal B and HER2+ into Luminal, and TNBC and Claudin-low into Basal. Data was further subset to Luminal and Basal subtypes only (62 out of 67 cell lines), resulting in an overall composition of 33 Luminal and 29 Basal cell lines. Finally, response was aggregated across cell lines of the same subtype/lineage grouping by marker, treatment, timepoint and lineage itself, and displayed as mean (solid lines) and standard error of the mean (SEM, shaded regions) across timepoints shared by at least 4 cell lines.

#### Mass spectrometry and multiplexed tumour imaging

Mammary tumours were generated by injecting tumour organoids under the nipple into the mammary fat pad. Vehicle or 130 mg/kg Capivasertib was administered twice daily by oral gavage for 4 days, followed by 3 days off. Upon reaching the experimental endpoint, mice were sacrificed and tumours snap frozen using liquid nitrogen immediately after resection. The tissues were embedded in a hydroxypropyl methylcellulose (HPMC)/polyvinylpyrrolidone (PVP) hydrogel as previously described (*57*). Sectioning was performed on a CM3050 S cryostat (Leica Biosystems, Nussloch, Germany) at a section thickness of 10 µm and the tissue sections were immediately thaw mounted and dried under a stream of nitrogen and sealed in vacuum pouches.

##### Mass spectrometry imaging

Tissue sections for DESI (Desorption Electrospray Ionization)-MSI were thaw-mounted onto Superfrost microscope slides, while sections prepared for MALDI (Matrix-Assisted Laser Desorption Ionization)-MSI were thaw mounted onto conductive ITO coated slides). PVP (MW 360 kDa) and HPMC (viscosity 40–60 cP, 2% in H2O (20 C) were purchased from Merck (Darmstadt, Germany). Acetonitrol, water, isopentane and isopropyl alcohol were obtained from Fisher Scientific (Waltham, Massachusetts, USA).

DESI-MSI was perfomed for endogenous metabolite imaging. DESI-MSI analysis was performed on a Exploris 480 mass spectrometer (Thermo Scientific, Bremen, Germany) equipped with an automated two-dimensional-DESI ion source (Prosolia, Indianapolis, Indy, USA) operated in negative ion mode, covering the applicable mass range up to an m/z of 1000, with a nominal mass resolution of 70,000. The injection time was fixed to 150 ms. The spatial resolution used was 65 µm. A home-built Swagelok DESI sprayer was operated with a mixture of 95% methanol, 5% water delivered with a flow rate of 1.5 µL/min and nebulized with nitrogen at a backpressure of 6 bar. The resulting .raw files were converted into .mzML files using ProteoWizard msConvert (*58*) (V.3.0.4043) and subsequently compiled to an .imzML file (imzML converter (*59*) V.1.3). All subsequent data processing was performed in SCiLS Lab (V.2026a, Bruker Daltonik, Bremen, Germany).

MALDI-MSI was performed for Capivasertib distribution imaging. MALDI-MSI analysis was performed on a timsTOF flex instrument (Bruker Daltonik, Bremen, Germany) operated in positive detection mode. 2,5-Dihydroxybenzoic acid was prepared in 50:50 acetonitrile:water with 0.1% trifluoracetic acid at a concentration of 37.5mg/ml was used as an MALDI matrix. The matrix was spray deposited using an automated spray system (M3-Sprayer, HTX technologies, Chapel Hill, North Carolina, USA). The settings used were 100 µL/min flow rate, 8 passes, 75°C. MALDI experiments were performed with a spatial resolution of 50 µm and a mass range of 300-1000. A total of 200 laser shots were summed up per pixel to give the final spectra. Collision RF used was 1100 and transfer time 50. For all experiments the laser was operated with a repetition rate of 10 kHz. All raw data were directly uploaded and processed in SCiLS lab (V.2026a) software packages. All DESI and MALDI data and images were normalized to root mean square. For the MALDI data, unsupervised segmentation of the tissue images into regions was performed with bisecting K-means using the 250 features most frequently present across the image. For DESI data, tissue regions were classified using linear discriminant analysis using the 500 top features. The results were compared to overlayed H&E images to select regions from the MSI analysis which closely represented viable tumour regions in the H&E image. Average intensity values across the viable tumour regions were exported for downstream plotting and analysis.

##### Multiplexed tumour imaging

Slides were washed and incubated at 4°C for 10 minutes in solution of 2% H2O2 in methanol to block endogenous peroxidase. Slides were washed and then rinsed in 100% IMS and air dried before loading onto Leica Bond Rx platform. Antigen retrieval stripping steps between each antibody were performed with Epitope Retrieval Solution 1 (pH9) (Leica, AR9961) for 20 minutes at 95_°_C. Slides were then incubated with 0.1% BSA-PBST solution for 5 minutes at RT for protein blocking prior to each antibody.

Multiplex immunofluorescence was performed on a BondRx Leica Bond Rx, and antibodies were applied with Opal pairings in the following order: Ki67 (abcam, ab15580) at 1:1500 dilution with Opal 570 (Akoya, FP1488001KT) at 1:300 dilution, Cytokeratin-14 (abcam, ab7800) at 1:500 dilution with Opal 520 (Akoya, FP1487001KT) at 1:300 dilution, pS6 (CST, 2211S) at 1:400 dilution with Opal 620 (Akoya, FP1495001KT) at 1:300 dilution, Laminin (Sigma, L9393) at 1:100 dilution with Opal 780 (Akoya, FP1501001KT) at 1:100 and 1:25 dilution. Anti-rabbit Polymer (Leica, RE7260-CE, RTU) was used as secondary for antibodies raised in rabbit and rabbit anti-mouse Post Primary (Leica, RE7260-CE, RTU) and anti-rabbit Polymer (Leica, RE7260-CE, RTU) were used as secondary for antibodies raised in mouse. Antibodies were diluted in 1xPlus automation amplification antibody diluent (Akoya, FP1609) except for pS6, which was diluted in 1X Antibody Diluent/Block (Akoya, ARD1001EA). Slides were counterstained with DAPI (Thermo Scientific, 62248) 1:2500 and mounted with Prolong Gold Antifade reagent (Invitrogen, P36934). Slides were scanned using the PhenoImager HT (formerly Vectra Polaris) using PhenoImager HT 2.0 software.

### Supplemental Text

Live imaging approaches are well-suited to recording longitudinal data before and after therapeutic challenge and the spatial context of cells, with biosensors being deployed to reveal information about cell signalling and state (*60–67*). The advent of light-sheet imaging has transformed our ability to image three-dimensional models that closely recapitulate the biology of tumours, such as organoids and tissue explants. In particular, the development of light-sheet fluorescence microscope systems is beginning to enable large numbers of 3D structures to be imaged in parallel (*68–70*).

We leveraged a custom-built dual-view oblique plane microscope (dOPM) to perform multi-dimensional live-cell imaging compatible with a multi-well plate format (Fig. S3A). GCRC1915 organoids were transduced using a piggyBac system to express a modified segment of the AKT substrate FOXO1 (FOXO1_10A3D_-mRuby2) that accumulates in the cytoplasm when phosphorylated (*71*), a glucose biosensor that reports on glycolysis (*63, 72*), and a nuclear label for image processing purposes (*73*) (Fig. S2B & C). A bespoke pipeline was set up to analyse AKT activity and glucose levels in every cell of every organoid (Fig. S3D). Individual organoids were assigned a unique identity, and all cell metrics were linked to the organoid identity (Fig. S3E). Within the organoids, nuclei were segmented in 3D based on the H2B-miRFP670nano signal. This was used to generate both a nuclear mask and a perinuclear mask, encompassing a one-micron thick shell immediately outside the nucleus. This enabled us to determine the number of nuclei per organoid and derive an organoid growth metric. AKT activity was determined by dividing the FOXO1_10A3D_-mRuby2 signal in the perinuclear mask by the sum of the nuclear mask and the perinuclear mask, with higher values indicating greater AKT activity. Intracellular glucose levels were determined by ratiometric analysis of the Citrine sensitised emission and eCFP signals. Fig. S2E shows 50 representative DMSO and Capivasertib-treated organoids from one of 4 experimental replicates. Thus, we established methodology enabling direct *in situ* assessment of drug efficacy, the coupling of AKT to glycolysis, and organoid growth at scale.

As described above, metrics were determined for every segmented nucleus, with each cell being assigned to a specific organoid. Control experiments using Koningic acid and glucose-free media confirmed the fidelity of the glucose sensor, and additionally revealed an up-regulation of AKT activity when glucose was absent (Fig. S4A). Principal Component Analysis (PCA) incorporating both biosensor and morphological metrics were used to explore the diversity of cell states in the organoids. Fig. S4C shows changing cell metrics upon treatment. Cells treated with Koningic acid showed uniform changes. In contrast, Capivasertib-treated cells showed divergent responses with two distinct populations emerging at 96 hours. A similar divergence was observed when the response to Capivasertib was analysed at the organoid level, suggesting the majority of cells within an organoid respond in a similar manner to the drug (Fig. S4D). Fig. S4E confirms that cell states in any given organoid cluster together 96 hours after drug treatment. We next interrogated the functional difference between organoids, specifically their growth rate. Overlaying the growth rates indicated that the cluster containing the majority of the control organoids had a high growth rate, while the Capivasertib-treated organoids were split between the high growth rate cluster and a low growth rate cluster, thus confirming the analysis in Fig. 1 (Fig. S4F). Thus, our imaging platform reveals divergence in response to AKT inhibition at the organoid level.

**Figure S1.**
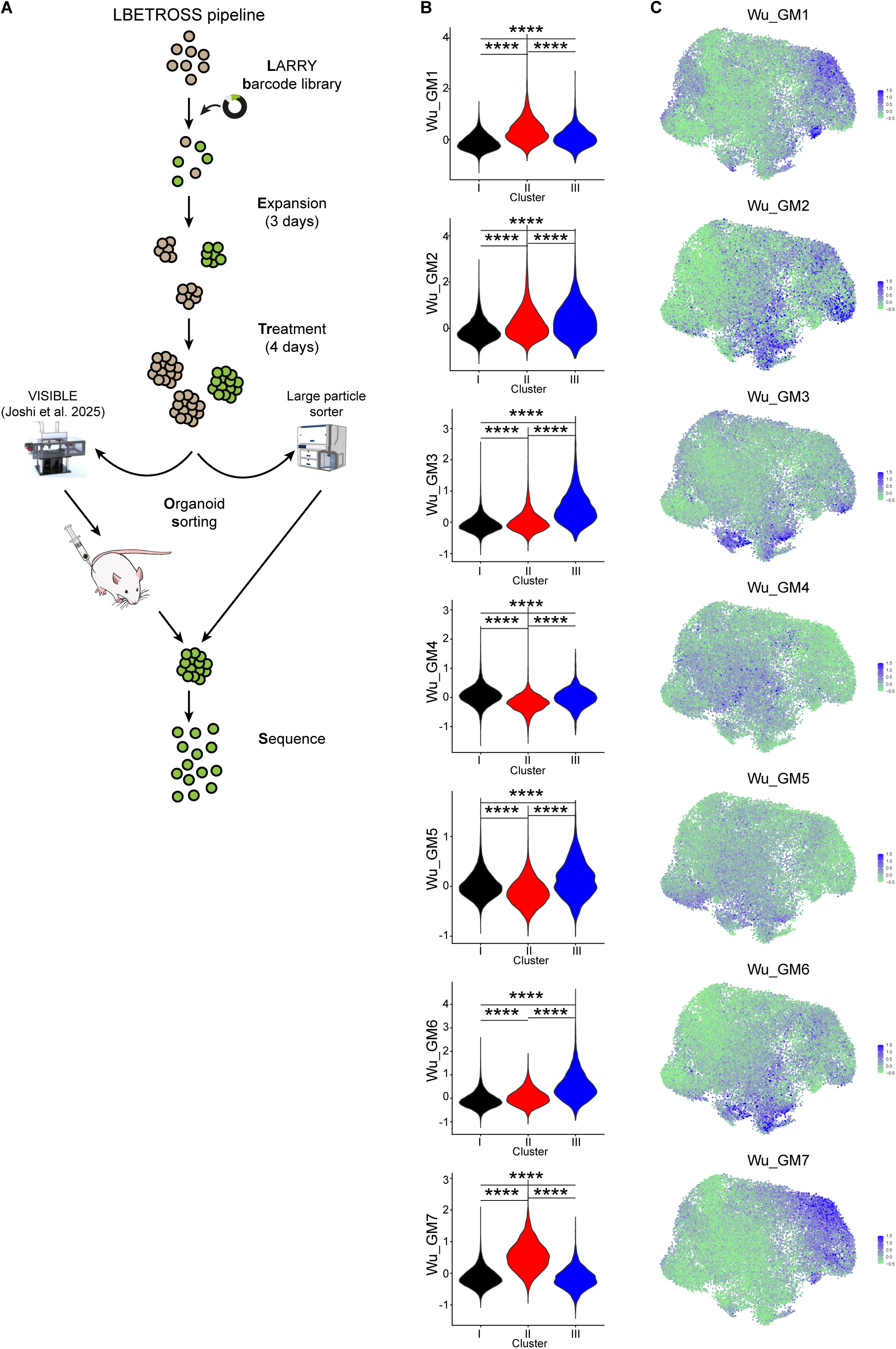
Gene module expression in a PDXOX model of triple negative breast cancer. **(A)** Schematic of experimental steps taken to prepare single-cell RNAseq datasets **(B)** Violin plots depicting the gene module scores as defined by Wu et al. for cells within each cluster of the LARRY labelled PDXOX dataset. **** p < 0.0001 (Wilcoxon test). **(C)** UMAP representation of GCRC1915 patient-derived xenograft single-cell RNA sequencing datasets with colours indicating expression of gene modules.

**Figure S2.**
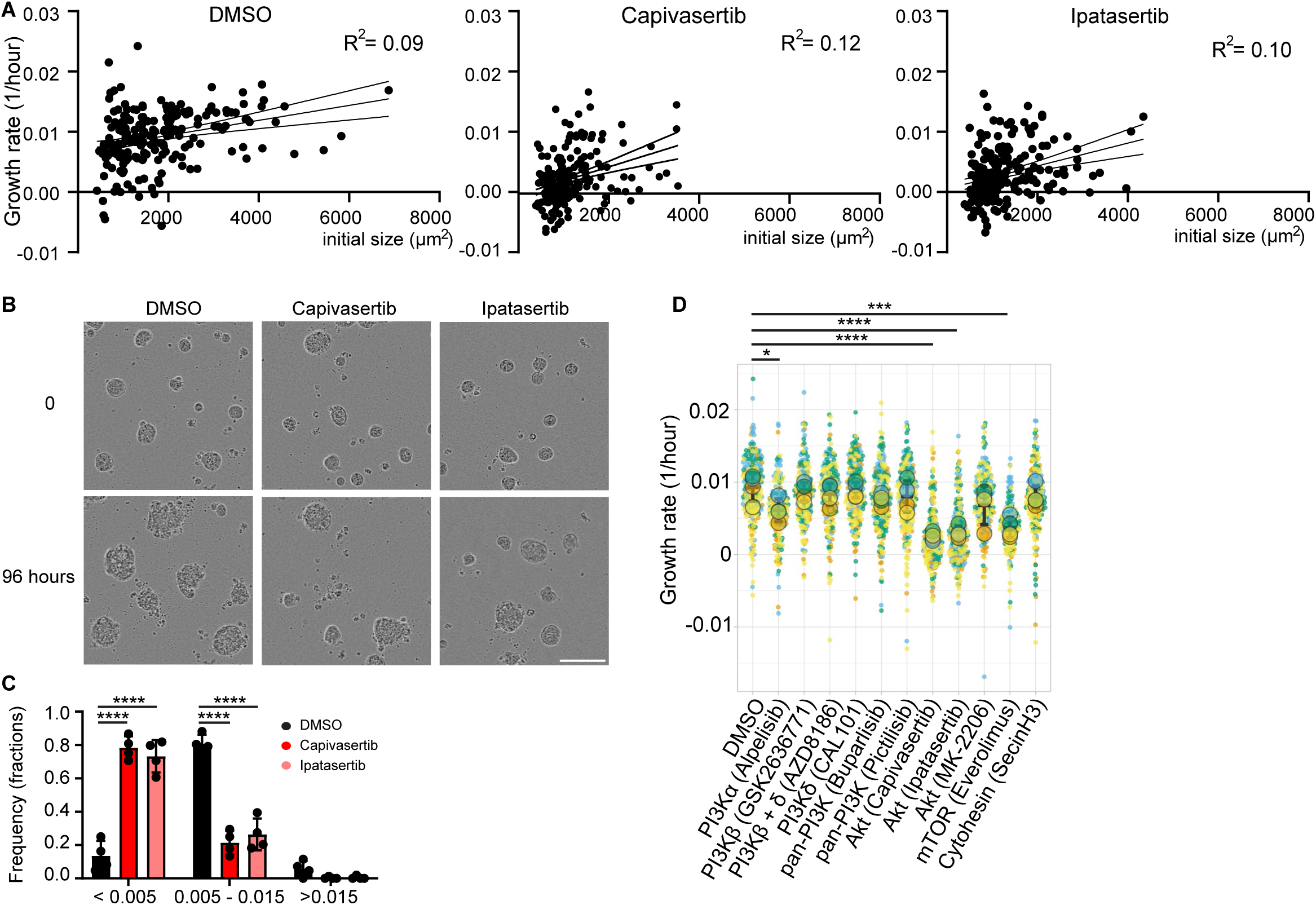
PDXO sensitivity to PI3K/Akt/mTOR pathway inhibitors. **(A)** Growth rate of individual organoids treated with DMSO, 1 μM Capivasertib or 1 μM Ipatasertib are plotted as a function of their initial size. **(B)** Representative brightfield micrographs of GCRC1915 PDXOs. Scale, 50 μm. **(C)** Relative frequency histogram of organoid growth rates for each indicated condition. **(D)** Growth rates of organoids treated with indicated drug conditions for 4 days. Data are represented as Superplots where dots represent organoids (n=2263), circles represent experimental means (n=4) and colours indicate experiment identity. * p < 0.05 ***p < 0.001 **** p < 0.0001 (Holm-Šídák’s multiple comparison test).

**Figure S3.**
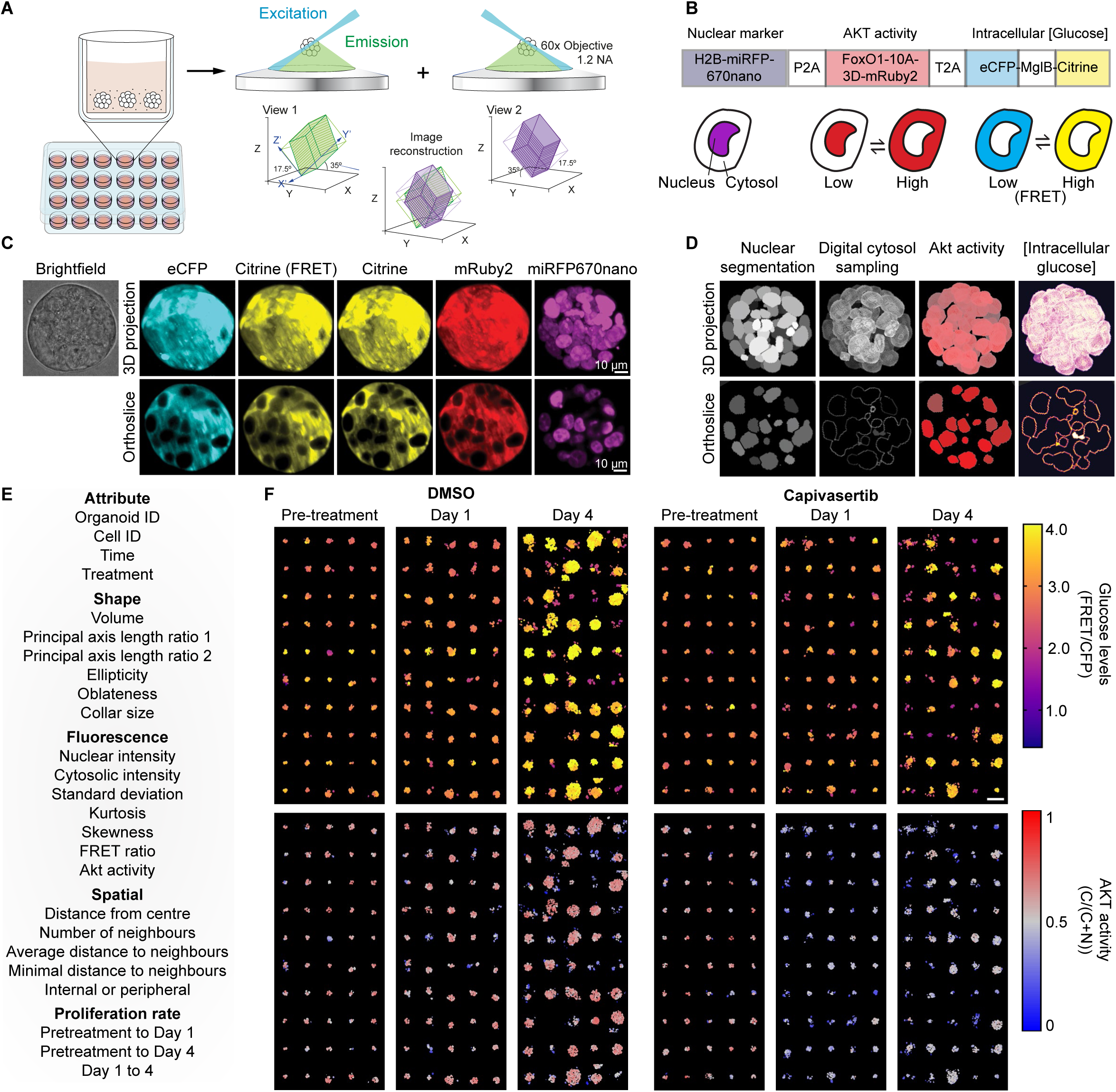
dOPM enables single-cell and whole organoid metrics of PI3K pathway activity. **(A)** Schematic of organoids relative to the objective of the dOPM. **(B)** Schematic of the polycistronic construct encoding a nuclear marker, Akt activity reporter and intracellular glucose biosensor separated by P2A or T2A self-cleaving peptide sequences. **(C)** Representative brightfield or fluorescent OPM micrographs of biosensor GCRC1915 PDXO organoids Scale, 10 μm **(D)** Representative masks of quantitative metrics derived from fluorescent construct. **(E)** Digital montage of maximum project images of pseudo-coloured micrographs representing intracellular glucose levels (left) and Akt activity (right) from 50 DMSO (top) and 50 Capivasertib (bottom) treated organoids immediately prior to, 1 day or 4 days after treatment. Scale, 100 μm.

**Fig. S4.**
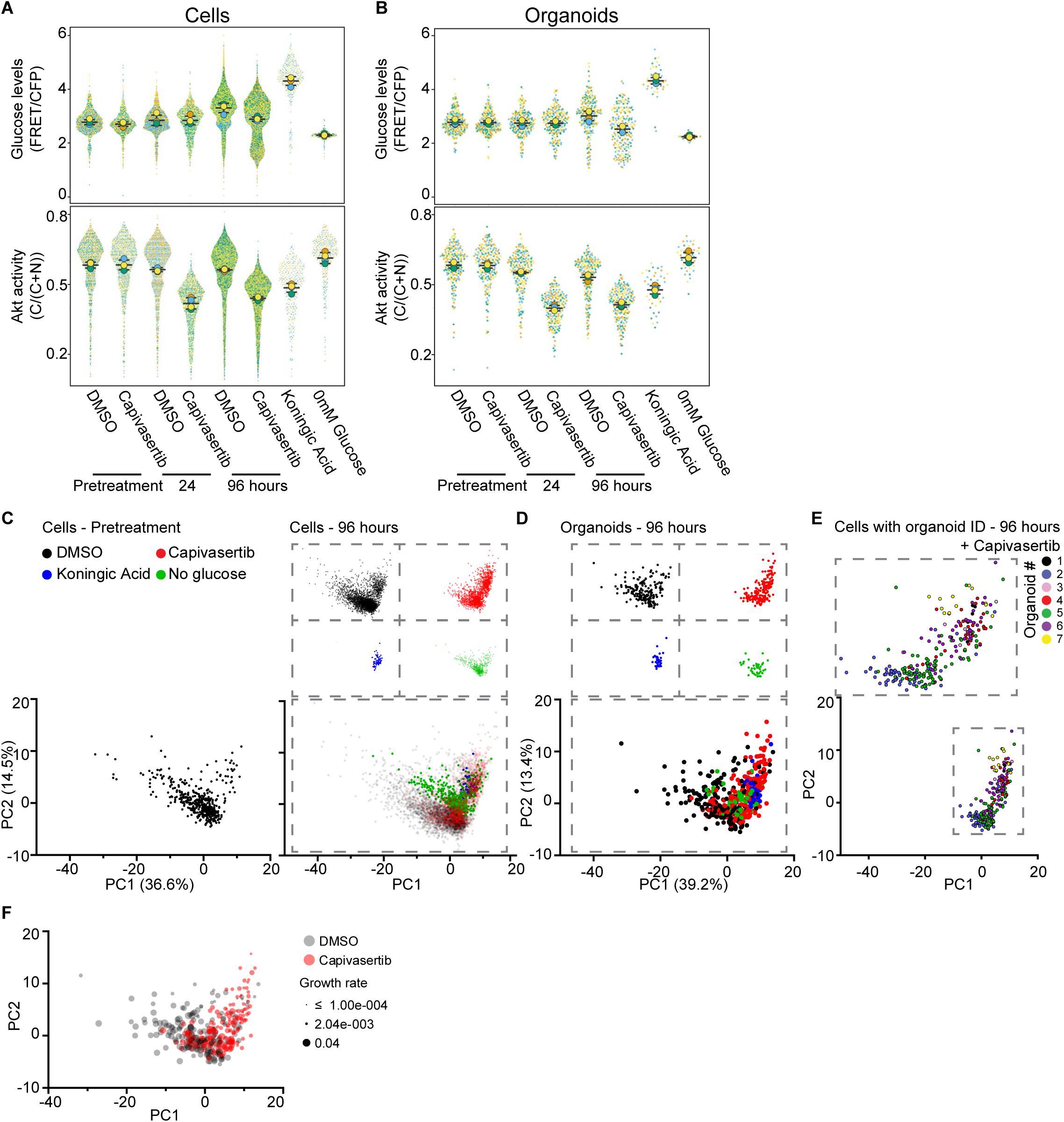
Overview of cellular and whole organoids metrics of PI3K pathway activity. (**A**) Glucose levels and AKT activity of cells treated with indicated drug conditions. Data are represented as Superplots where dots represent cells (n=22 044), circles represent experimental means (n=4) and colours indicate experiment identity. **(B)** Values of median glucose levels and Akt activity of organoids treated with indicated drug conditions. Data are represented as Superplots where dots represent median organoid levels (n=679), circles represent experimental means (n=4) and colours indicate experiment identity. (**C, D. and E)** Principal component analysis of biosensor-derived metrics from **(C)** single cells and (**D)** median values of whole organoids (480 organoids, n=4 experimental replicates). Regions indicated in the bottom-right grey dashed box are shown above for individual treatments. **(E)** Principal component analysis of biosensor-derived metrics from single cells of 7 organoids. Colours denote organoid identity. **(F)** Principal component analysis of biosensor-derived metrics from median values of whole organoids. Colour denotes treatment condition (DMSO – black & 1 μM Capivasertib red) and dot size indicates organoid growth rate between 0 and 96 hours.

**Fig. S5.**
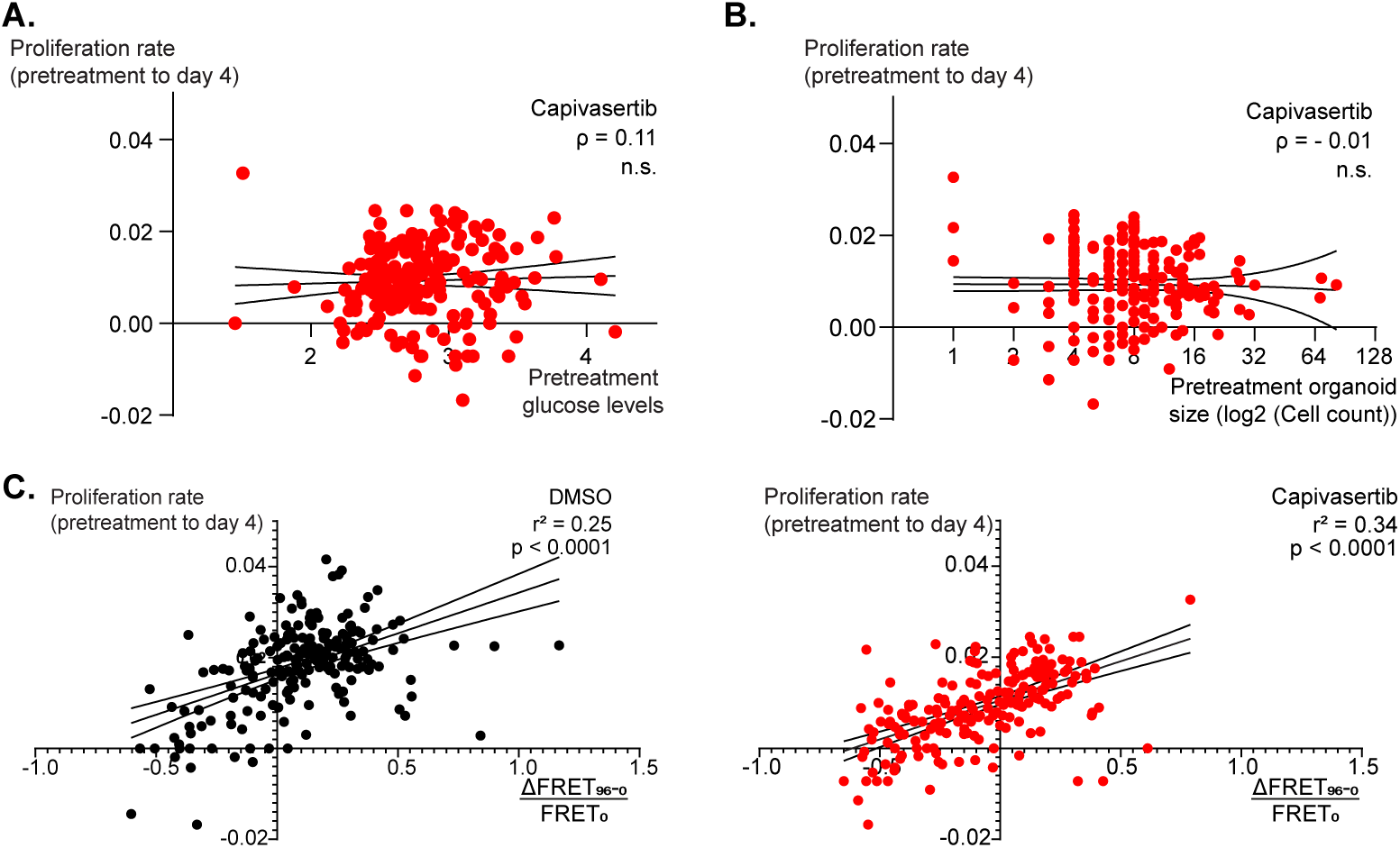
Comparison of proliferation with selected dOPM derived metrics. **(A)** Plotted proliferation rates of organoids with 1 μM Capivasertib against pretreatment glucose levels. **(B)** Plotted proliferation rates of organoids with 1 μM Capivasertib against pretreatment organoid size. **(C)** Plotted growth rates of organoids with DMSO or 1 μM Capivasertib against the change in FRET values from the glucose biosensor relative to the initial values. Spearman’ correlation * p < 0.05 two-tailed.

**Fig. S6.**
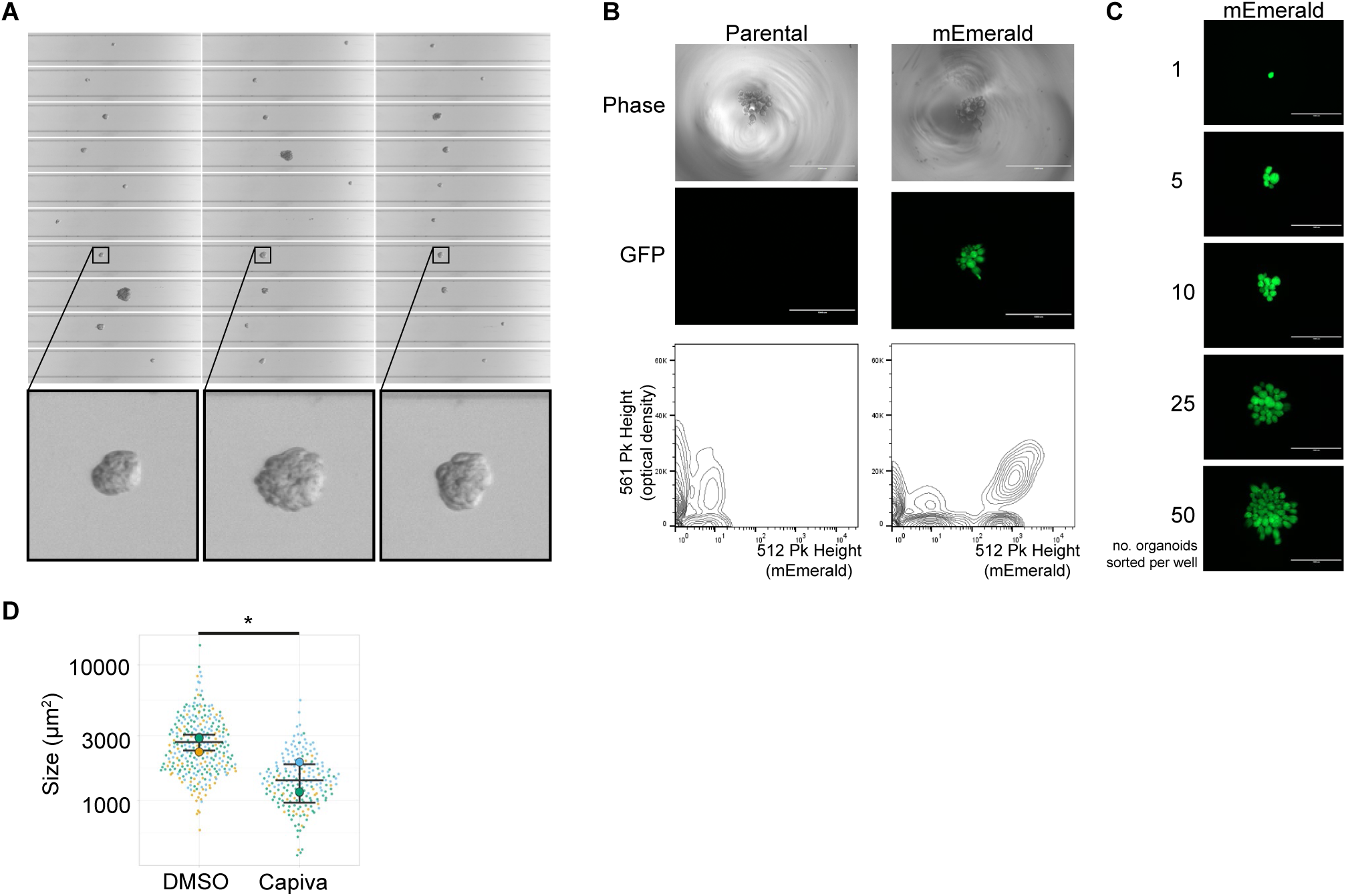
Sorting whole fluorescent organoids. **(A)** Representative images of organoids passing through the flow cell. **(B)** Brightfield and fluorescent micrograph as well as corresponding contour plost of parental or mEmerald expressing organoids sorted into a ultra-low adherent v-bottom well. **(C)** Fluorescent micrographs of specific numbers of sorted mEmerald+ organoids. **(D)** Values of organoid size derived from flow cell micrographs of sorted organoids treated with either DMSO or 1 μM Capivasertib. Data are represented as Superplots where dots represent organoids (n=490), circles represent experimental means (n=3) and colours indicate experiment identity. * p < 0.05 Unpaired two-tailed t-test.

**Fig. S7.**
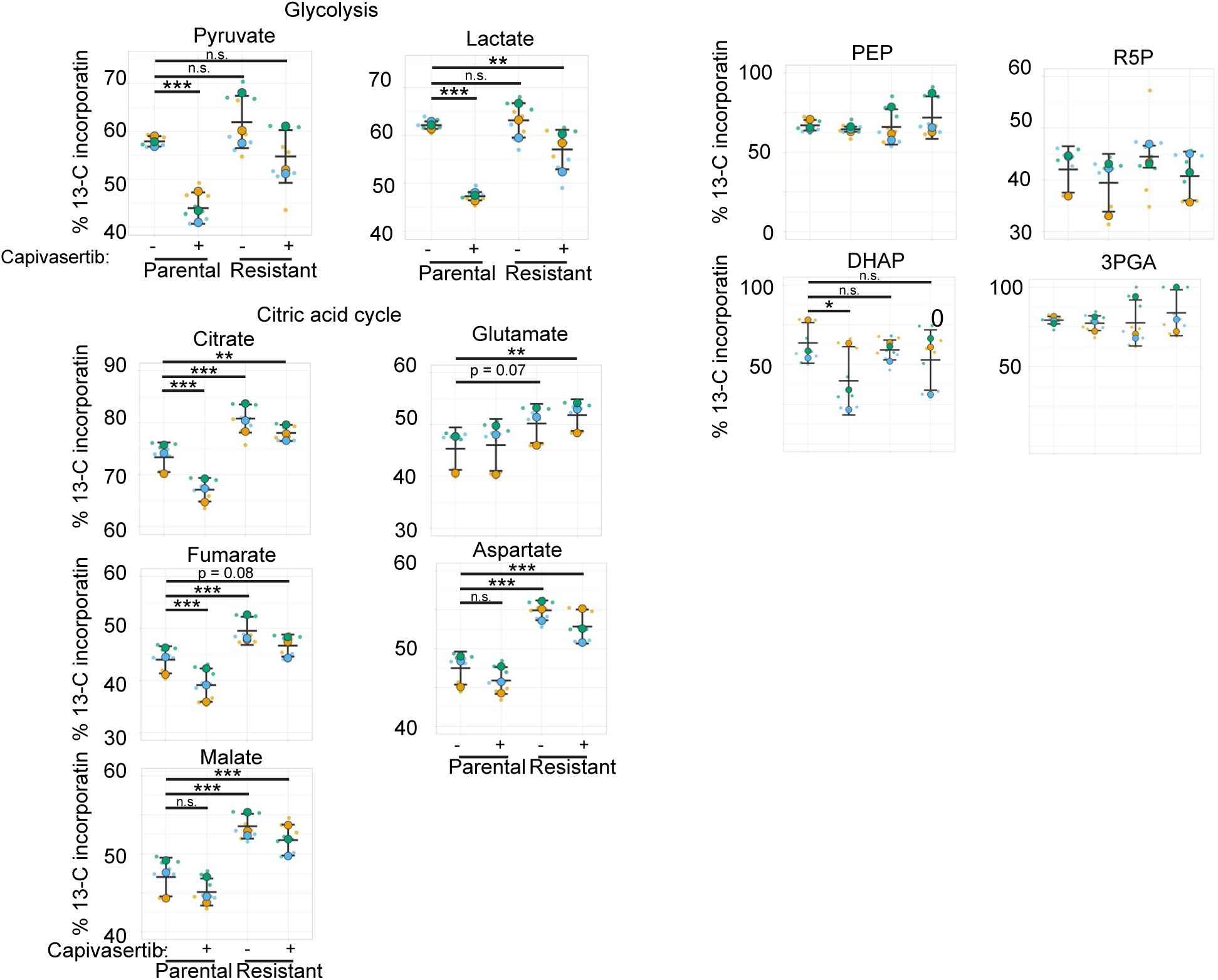
Metabolomic analysis of Capivasertib resistant organoids. % 13-C-Glucose incorporation of selected glycolytic and TCA cycle intermediates in parental or Capivasertib resistant organoids treated with either DMSO or 1 μM Capivasertib. Data are represented as Superplots where dots represent technical replicates (8 or 9), circles represent biological replicate means (n=3) and colours indicate biological replicate identity. * p < 0.05 ** p<0.01 *** p <0.001.

**Fig. S8.**
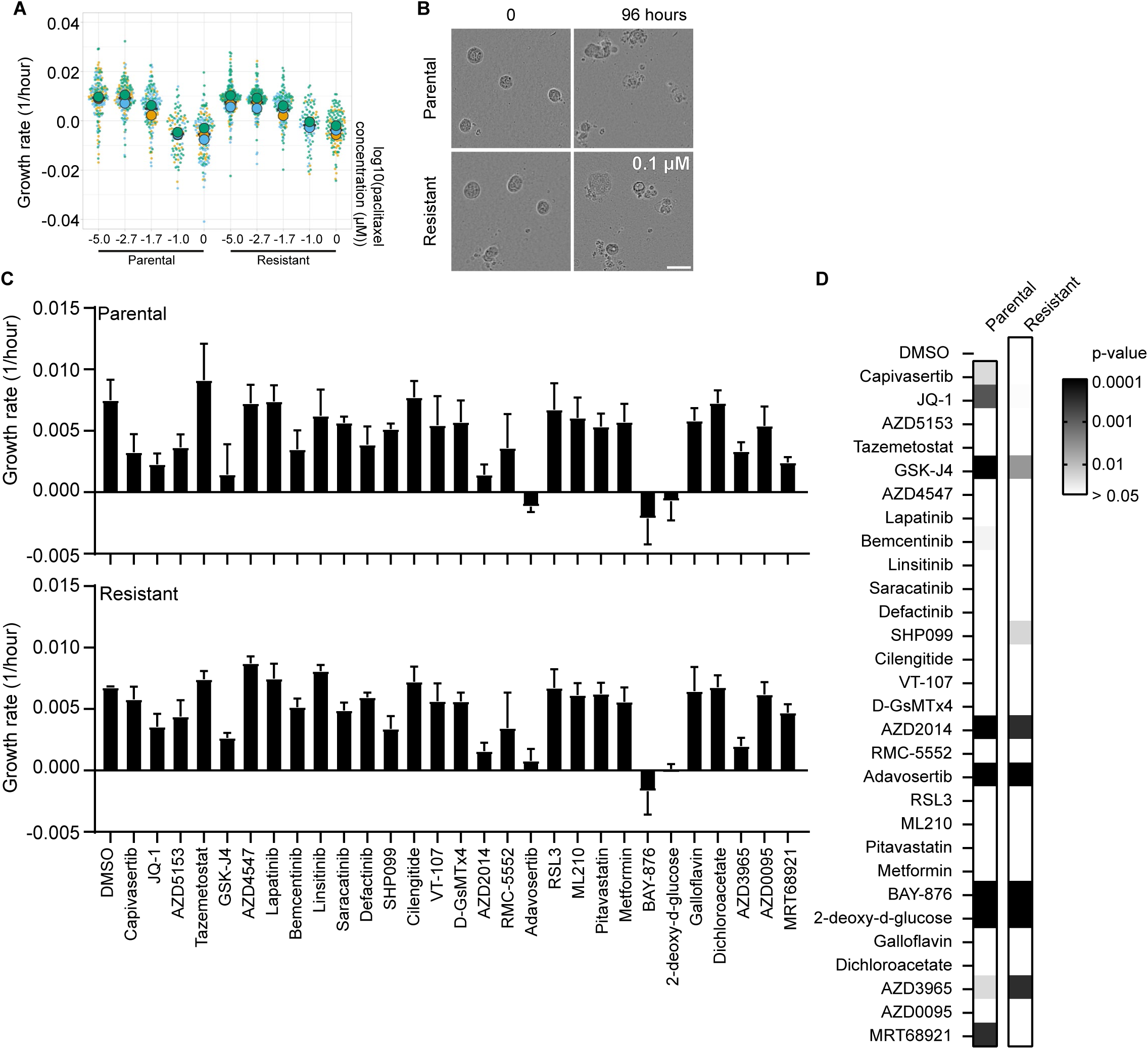
Drug sensitivity of parental and Capivasertib resistant organoids. **(A)** Growth rates of parental or resistant organoids treated with indicated concentration of paclitaxel for 4 days. Data are represented as Superplots where dots represent organoids (n=1444), circles represent experimental means (n=3) and colours indicate experiment identity. **(B)** Representative brightfield micrographs of GCRC1915 PDXOs. Scale, 50 μm. **(C)** Mean growth rates of parental or Capivasertib-resistant organoids treated with the indicated drugs. (n=3) **(D)** Heatmap of statistical analysis of growth rates compared to DMSO treated parental controls using Dunnett’s multiple comparison test.

**Fig. S9.**
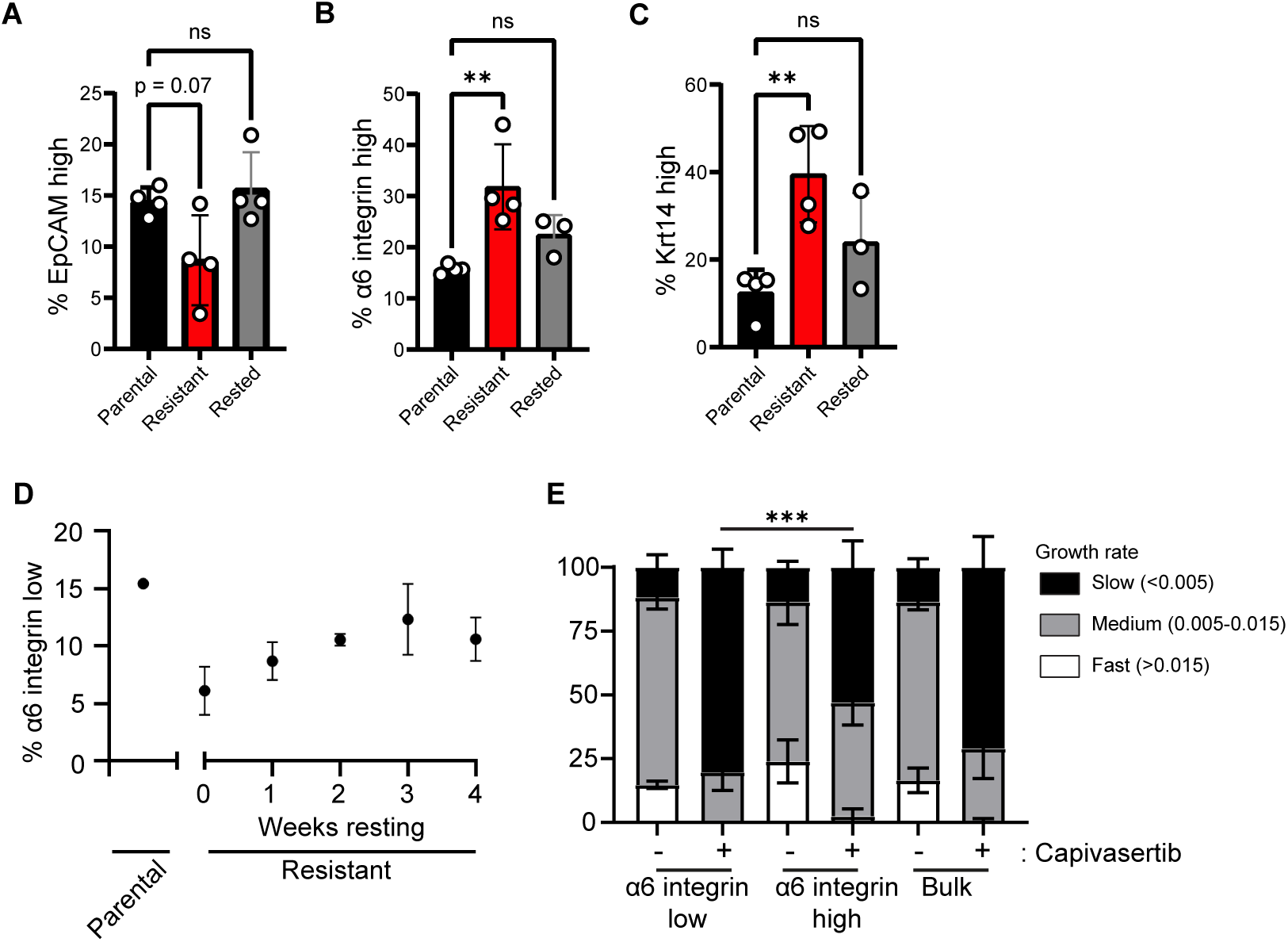
Capivasertib-resistant organoids exhibit reversible basal-like features. **(A, B & C)** Barplot of flow cytometry analysis of organoid-derived cells stained for indicated markers. (n=3 or 4) **(D)** Flow cytometry analysis of a α6 integrin low subpopulation in DMSO or Capivasertib-resistant organoids rested for the indicated periods. (n=3) **(E)** Relative fractions of organoid growth rates derived from α6 integrin low, high or bulk-sorted cells. After 3 days of growth, organoids were imaged for 4 days. (n=3) ** p<0.01 *** p<0.001 **** p<0.000. (Dunnett’s multiple comparison test)

**Fig. S10.**
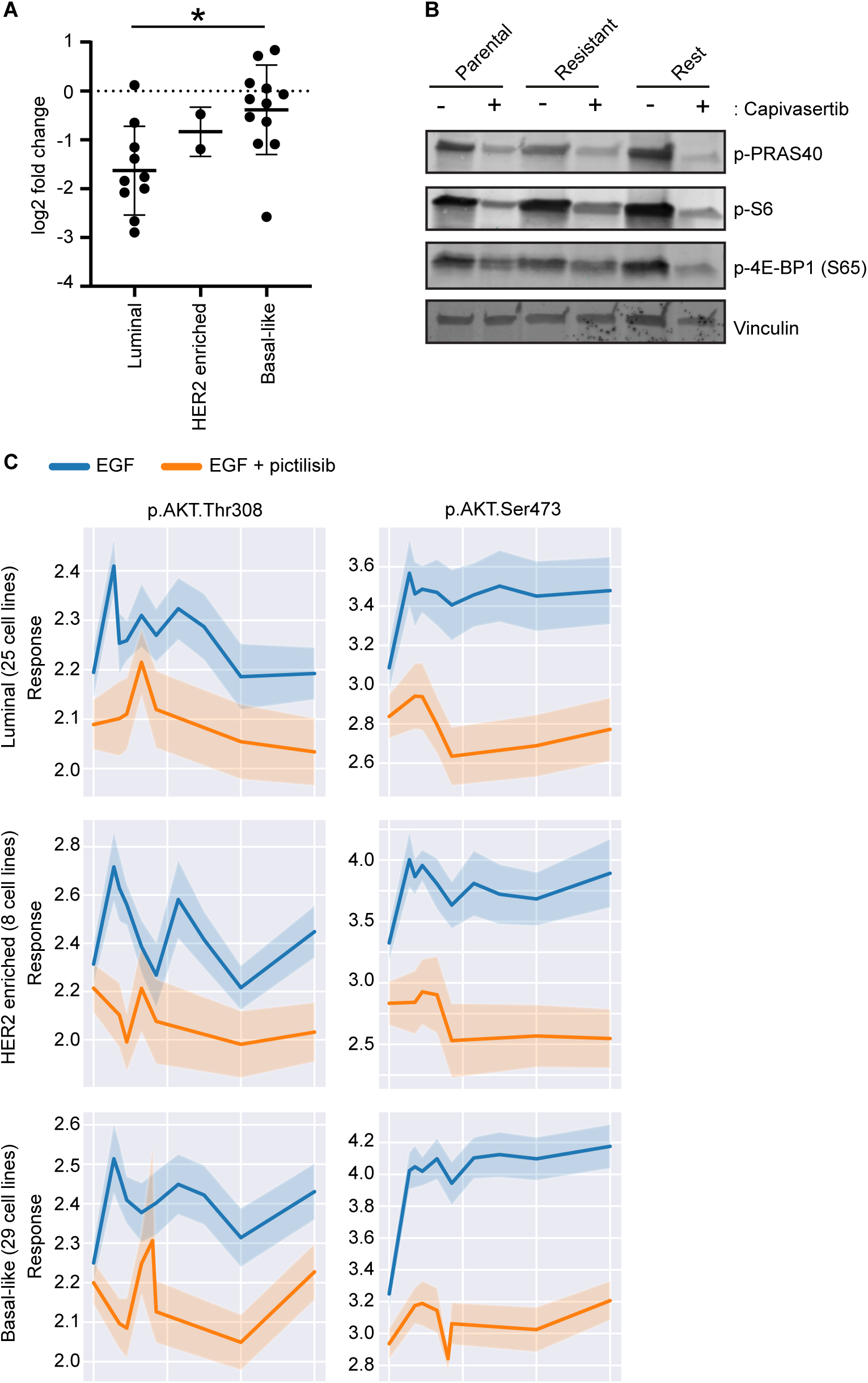
Analysis of breast cancer intrinsic subtype response to Capivasertib. **(A)** Comparison of breast cancer cell line viability separated according to intrinsic breast cancer subtype (downloaded from DepMap.org version PRISM Repurposing Public 24Q2). **(B)** Western blot from parental, resistant or rested GCRC1915 PDXOs treated with DMSO or 1 μM Capivasertib for 24 hours. **(C)** Comparison of CyTOF PI3K signalling data in breast cancer cell lines separated according to intrinsic breast cancer subtype.

**Fig. S11.**
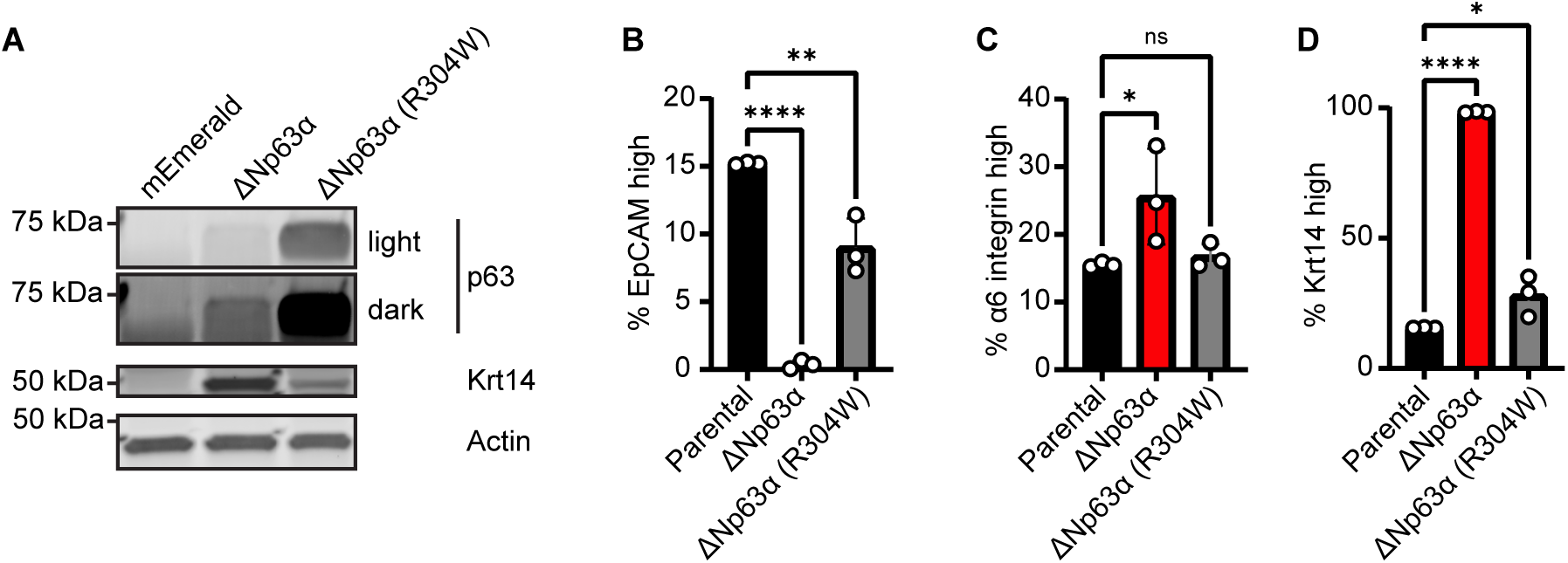
Characterisation of p63-overexpressing organoids. **(A)** Western blot of GCRC1915 PDXOs stably overexpressing the indicated constructs. **(B, C & D)** Barplot of flow cytometry analysis of organoid-derived cells stained for indicated markers. (n=3) ** p<0.01 *** p<0.001 **** p<0.000. (Dunnett’s multiple comparison test).

**Fig. S12.**
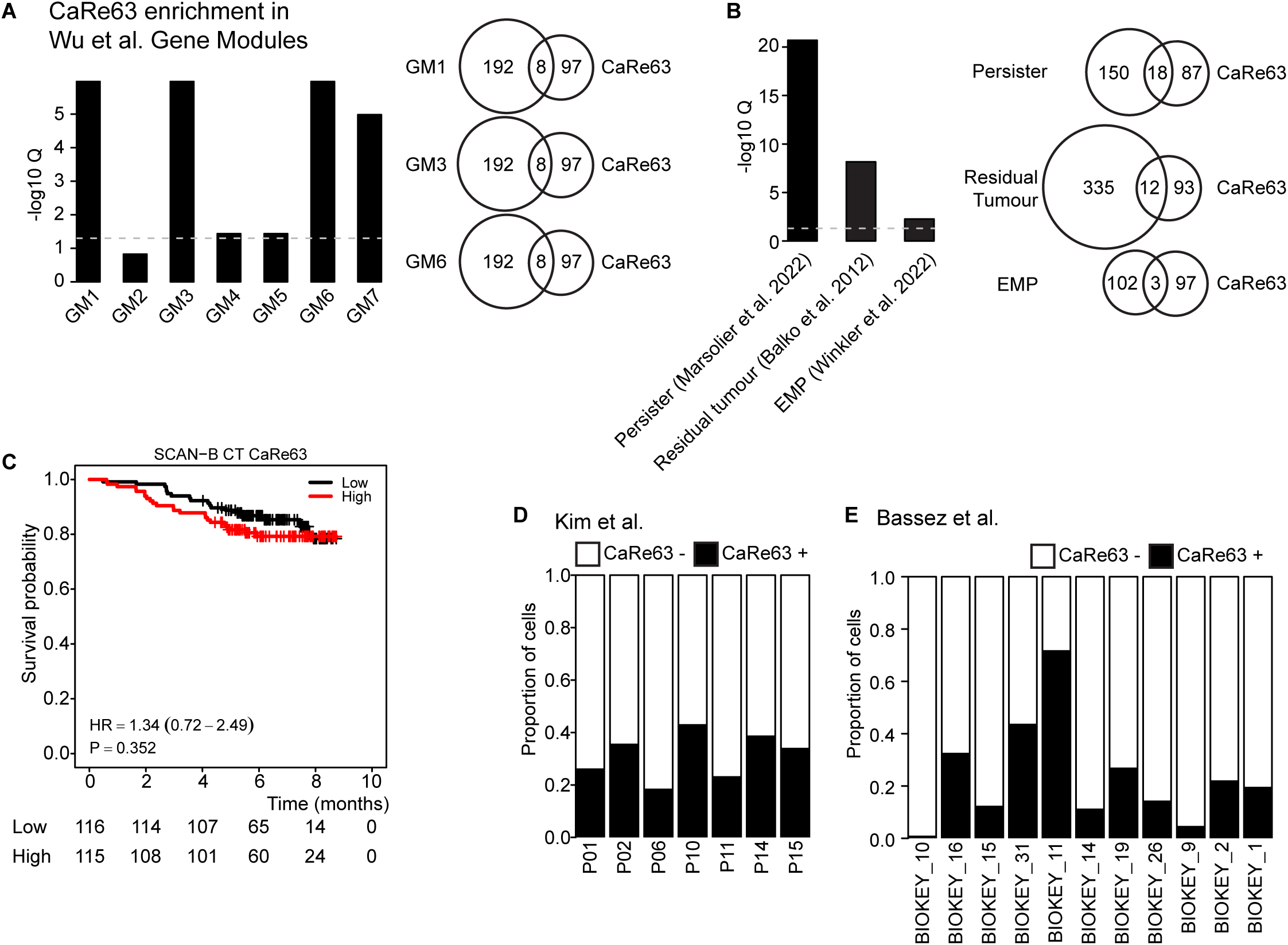
Analysis and comparison of CaRe63 in publicly available TNBC datasets. **(A)** Enrichment analysis of CaRe63 in gene modules defined in Wu et al. **(B)** Enrichment analysis of CaRe63 in chemoresistance-associated gene signatures. **(C)** Kaplan-Meier curve of patients in the SCAN-B cohort of TNBC cases treated with chemotherapy separated by low and high CaRe63 levels. **(D, E & F)** Proportion of CaRe63+ (GSVA > 0) cells in indicated patient and patient-derived xenograft single cell datasets.

**Fig. S13.**
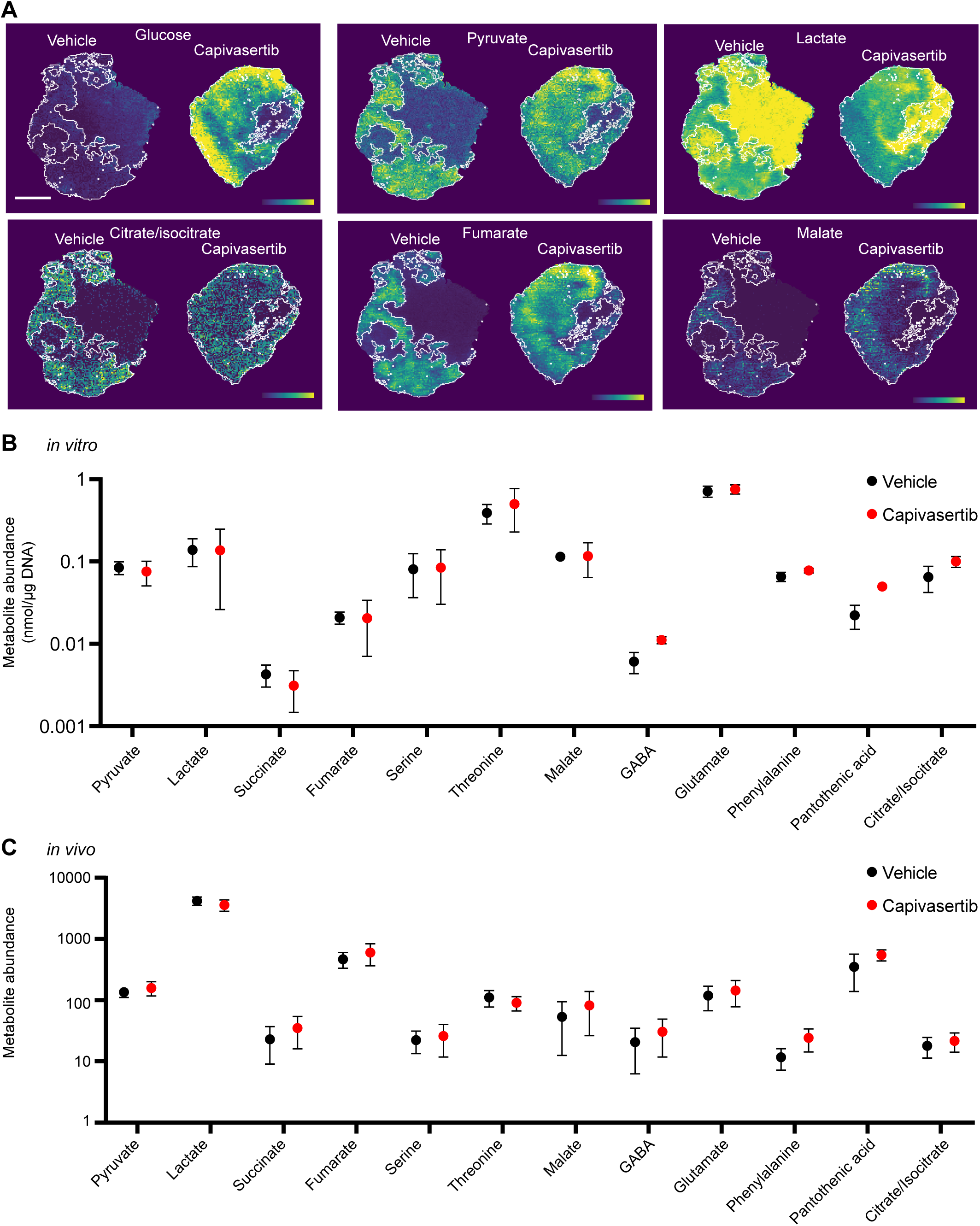
Analysis of metabolite abundance in Capivasertib-treated organoids and tumours. **(A)** Spatial distribution maps of metabolites in representative vehicle or Capivasertib treated tumours. Scale, 2 mm. **(B)** Bar plots of selected metabolite abundance in parental or Capivasertib-resistant organoids treated with either DMSO or 1 μM Capivasertib. (n=2 or 3) **(C)** Bar plots of selected metabolite abundance in GCRC1915 PDXOX derived tumours treated with either vehicle (n=7) or Capivasertib (n=6).

**Table S1.**

| CaRe63 |
| --- |
| ENSG00000104368 |
| ENSG00000165272 |
| ENSG00000145824 |
| ENSG00000120471 |
| ENSG00000181449 |
| ENSG00000125148 |
| ENSG00000129194 |
| ENSG00000136997 |
| ENSG00000171223 |
| ENSG00000034510 |
| ENSG00000153113 |
| ENSG00000068366 |
| ENSG00000114541 |
| ENSG00000107984 |
| ENSG00000101871 |
| ENSG00000114861 |
| ENSG00000285708 |
| ENSG00000156299 |
| ENSG00000099810 |
| ENSG00000109113 |
| ENSG00000156535 |
| ENSG00000114115 |
| ENSG00000075618 |
| ENSG00000144824 |
| ENSG00000132669 |
| ENSG00000134762 |
| ENSG00000138829 |
| ENSG00000166681 |
| ENSG00000003096 |
| ENSG00000198873 |
| ENSG00000154122 |
| ENSG00000122786 |
| ENSG00000076356 |
| ENSG00000177494 |
| ENSG00000168785 |
| ENSG00000175793 |
| ENSG00000189334 |
| ENSG00000134294 |
| ENSG00000120337 |
| ENSG00000119771 |
| ENSG00000101384 |
| ENSG00000188001 |
| ENSG00000105971 |
| ENSG00000182197 |
| ENSG00000165474 |
| ENSG00000105810 |
| ENSG00000089356 |
| ENSG00000125968 |
| ENSG00000117318 |
| ENSG00000159166 |
| ENSG00000159388 |
| ENSG00000118257 |
| ENSG00000143479 |
| ENSG00000149256 |
| ENSG00000132470 |
| ENSG00000058085 |
| ENSG00000106571 |
| ENSG00000131016 |
| ENSG00000170786 |
| ENSG00000213626 |
| ENSG00000111817 |
| ENSG00000100739 |
| ENSG00000100234 |
| ENSG00000096696 |
| ENSG00000175984 |
| ENSG00000154080 |
| ENSG00000172292 |
| ENSG00000151914 |
| ENSG00000164171 |
| ENSG00000206075 |
| ENSG00000139318 |
| ENSG00000170873 |
| ENSG00000163513 |
| ENSG00000188042 |
| ENSG00000184254 |
| ENSG00000163032 |
| ENSG00000140937 |
| ENSG00000134757 |
| ENSG00000105976 |
| ENSG00000026508 |
| ENSG00000113070 |
| ENSG00000058404 |
| ENSG00000084207 |
| ENSG00000138795 |
| ENSG00000187720 |
| ENSG00000107438 |
| ENSG00000050405 |
| ENSG00000149269 |
| ENSG00000112378 |
| ENSG00000160789 |
| ENSG00000188158 |
| ENSG00000012232 |
| ENSG00000164932 |
| ENSG00000074410 |
| ENSG00000198729 |
| ENSG00000145555 |
| ENSG00000147889 |
| ENSG00000058668 |
| ENSG00000091136 |
| ENSG00000163629 |
| ENSG00000152104 |
| ENSG00000156463 |
| ENSG00000157827 |
| ENSG00000176463 |
| ENSG00000106399 |
| ENSG00000186847 |

